# Inflammatory Cytokines Regulate T-Cell Development from Blood Progenitor Cells in a Stage and Dose-Specifc Manner

**DOI:** 10.1101/2021.01.18.427186

**Authors:** John M. Edgar, Peter W. Zandstra

## Abstract

T-cell development from hematopoietic stem and progenitor cells (HSPCs) is tightly regulated through Notch pathway activation by the Notch ligands Delta-like (DL) 1 and 4 and Jagged-2. Other molecules, such as stem cell factor (SCF), FMS-like tyrosine kinase 3 ligand (Flt3L) and interleukin (IL)-7, play a supportive role in regulating the survival, differentiation, and proliferation of developing progenitor (pro)T-cells. Numerous other signaling molecules are known to instruct T-lineage development *in vivo*, but little work has been done to optimize their use for T-cell production *in vitro*. Using a defined T-lineage differentiation assay consisting of plates coated with the Notch ligand DL4 and adhesion molecule VCAM-1, we performed a cytokine screen that identified IL-3 and tumor necrosis factor α (TNFα) as enhancers of proT-cell differentiation and expansion. Mechanistically, we found that TNFα induced T-lineage differentiation through the positive regulation of T-lineage genes *GATA3, TCF7*, and *BCL11b*. TNFα also synergized with IL-3 to induce proliferation by upregulating the expression of the IL-3 receptor on CD34^+^ HSPCs, yielding 753.2 (532.4-1026.9; 5-95 percentile)-fold expansion of total cells after 14 days compared to 8.9 (4.3-21.5)-fold expansion in conditions without IL-3 and TNFα. We then optimized cytokine concentrations for T-cell maturation. Focusing on T-cell maturation, we used quantitative models to optimize dynamically changing cytokine requirements and used these to construct a three-stage assay for generating CD3^+^CD4^+^CD8^+^ and CD3^+^CD4^−^CD8^+^ T-cells. Our work provides new insight into T-cell development and a robust *in vitro* assay for generating T-cells to enable clinical therapies for treating cancer and immune disorders.

## INTRODUCTION

T-cell immunotherapies using cancer antigen-specific T-cell receptors (TCRs) or chimeric antigen receptors (CARs) have emerged as a potent treatment option for diseases such as B-cell acute lymphoblastic leukemia (B-ALL) and diffuse large B-cell lymphoma^1^. These therapies collect peripheral T-cells from patients and expand and transduce them with a CAR before returning the T-cells back to the patient. While effective, the reliance on patient-derived cells is a limitation, as patients undergoing chemotherapy may not have a suitable number of cells for CAR T-cell therapy. Additionally, *in vitro* expansion can lead to T-cell exhaustion, limiting long-term efficacy^2^. The generation of T-cells from stem cells *in vitro* could provide an alternative source of cells for immunotherapies. Stem cells could be genetically engineered using CRISPR/Cas9 to target a CAR or antigen-specific TCR to the TCR α constant (TRAC) locus then differentiated to the T-lineage^3,4^. However, current methods for generating T-cells *in vitro* suffer from low efficiency and often produce mixed populations of T, natural killer (NK) and myeloid lineage cells. To generate T-cells in sufficient quantities for therapies, new strategies are needed to enhance overall cell yield while maintaining a relatively pure population of T-cells.

T-cell development *in vivo* occurs in the thymus. Hematopoietic progenitor cells migrate from bone marrow to the thymus where they initiate T-lineage development. Upon entry into the thymus, progenitors quickly upregulate CD7 and are defined as CD4/8 double negative (DN) CD34^+^CD7^+^(CD5^−^) proT1^5^. Throughout T-lineage specification, cells develop to become CD34^+^CD7^+^CD5^+^ proT2. Here, recombination of the TCRβ locus is initiated and is associated with T-lineage commitment. Cells begin to express CD4, passing through a CD4^+^ immature single positive (CDISP) stage before becoming CD4/8 double positive (DP). TCRαβ rearrangement are completed and cells undergo positive and negative selection. Finally, cells lose expression of either CD4 or CD8 depending on the specificity of the TCRαβ for a particular human leukocyte antigen (HLA) allele and enter the periphery as CD4^+^CD8^−^ or CD4^−^CD8^+^ single positive (SP) T-cells^6^.

T-lineage development is driven by Notch1 pathway activation through exposure to ligands Delta-like (DL) 1 and 4 and Jagged-2 expressed on the surface of thymic epithelial cells^7,8^. The protein vascular cell adhesion molecule 1 (VCAM-1) is co-expressed with Notch ligands on

TECs surface and facilitates migration through the thymus and continuous exposure to Notch ligands^9^. Other molecules present in the thymus, including stem cell factor (SCF) and interleukin (IL)-7, supplement Notch1 signaling to support T-cell survival, differentiation, and proliferation. Many other signaling molecules are also present in the thymus, including IL-1α/β, IL-3, IL-6, IL-12, and tumor necrosis factor α (TNFα)^10–12^, although their functions in human T-cell development are not as well understood.

A number of methods exist for generating T-cells *in vitro*^13–15^. One of the most successful was by co-culturing hematopoietic stem and progenitor cells (HSPCs) with mouse OP9 bone marrow stromal cells that ectopically express the Notch ligand DL1 or DL4 (OP9-DL)^14^. This system is robust and able to support development throughout proT and DP stages to generates CD8SP T-cells^16,17^. Other approaches have replaced OP9-DL with recombinant DL4-Fc fusion proteins immobilized to surfaces or microbeads^18,19^. Both serum and serum-free versions have been described with mixed success, and these are often limited to the early stages of T-lineage development.

We have previously described a serum-free system for generating proT-cells from human umbilical cord blood (UCB)-derived CD34^+^ HSPCs^20^. Called the DL4+VCAM-1 system, it uses plate-bound DL4 and VCAM-1 along with the recombinant cytokines SCF, FMS-like tyrosine kinase 3 ligand (Flt3L), IL-7, and thrombopoietin (TPO) to facilitate proT-cell development. While it generates proT-cells with similar efficiency to OP9-DL co-cultures, it provides only the minimum essential components necessary to do so. We hypothesized that incorporating additional thymus-associated soluble cytokines into DL4+VCAM-1 culture would enhance T-lineage differentiation and proliferation. Through a targeted cytokine screen, we identified IL-3 and TNFα as enhancers of T-lineage development. We found that TNFα accelerates T-lineage specification through interactions with the Notch1 pathway. Combining TNFα with IL-3 resulted in a significant expansion of proT-cells with minimal myeloid and NK cell contamination. We provide a method for quantitively predicting dynamic cytokine signaling requirements and show that TNFα switches from an enhancer to an inhibitor during T-cell development. Finally, we provide a three-stage protocol for generating DP and CD8SP T-cells in a system that is scalable and amenable for clinical translation.

## METHODS

### DL4 Production and DL4+VCAM-1 Plate Coating

Recombinant DL4-Fc fusion protein was purchased from Sino Biological or manufactured inhouse as previously described^20^. Tissue culture 96-well plates were coated with DL4-Fc and VCAM-1-Fc (R&D Systems) overnight at 4°C or for 2-4 hours at room temperature. To coat, DL4 and VCAM-1 were diluted to 15μg/ml and 2.5 μg/ml, respectively, in 50 μl of phosphate-buffered saline (PBS), resulting in a coating concentration of approximately 24 ng/mm^2^ of DL4 and 4 ng/mm^2^ of VCAM-1. For experiments with less DL4, the concentration of DL4 was adjusted accordingly while maintaining a 50 μl volume of PBS. Wells were washed once with PBS prior to seeding cells to remove unbound protein.

### Human CD34^+^ HSPC Enrichment from Umbilical Cord Blood

Umbilical cord blood (UCB) was collected from consenting donors at Mount Sinai Hospital, Toronto, Ontario or BC Children’s Hospital, Vancouver, British Columbia, in accordance with institutional research ethics board policies. CD34^+^ cells were isolated using the EasySep Human CD34^+^ Positive Selection Kit (Stemcell Technologies) to > 90% purity as previously described (Supplemental Figure 1)^20^. For certain experiments, enriched CD34^+^ cells were sorted into CD34^+^CD38^lo/-^ and CD34^+^CD38^+^ fractions using FACS Aria cytometer (Beckman Coulter).

### HSPC Culture on DL4+VCAM-1

For experiments 7 days or longer, HSPCs were seeded at 1000-4000 cells/well of a 96-well plate on DL4+VCAM-1-coated surfaces. Experiments shorter than 7 days (CFSE and qPCR) were seeded at 15000-25000 cells/well to provide enough cells for analysis. Cells were cultured in 100-200 μl of Iscove’s Modified Eagle Medium (IMDM) supplemented with 20% serum substitute (BIT 9500; Stemcell Technologies), 1 μg/ml low-density lipoprotein (LDL; Stemcell Technologies), 60 μM ascorbic acid (Sigma), 24 μM 2-mercaptoethanol (Sigma), and 1% penicillin-streptomycin (Invitrogen). The cytokines used in control condition were SCF, Flt3L, TPO, and IL-7 (all from R&D) at 100ng/ml for screening experiments (Figure 1) and 20ng/ml afterwards, unless otherwise mentioned. All other cytokines were purchased from R&D and used as described. For Notch pathway inhibition experiments (Figure 2), the γ-secretase inhibitor (2S)-N-[(3,5-Difluorophenyl)acetyl]-L-alanyl-2-phenyl]glycine 1,1-dimethylethyl ester (DAPT; R&D) or dimethyl sulfoxide (DMSO; Sigma) was added to media at the concentrations indicated.

**Figure 1.**
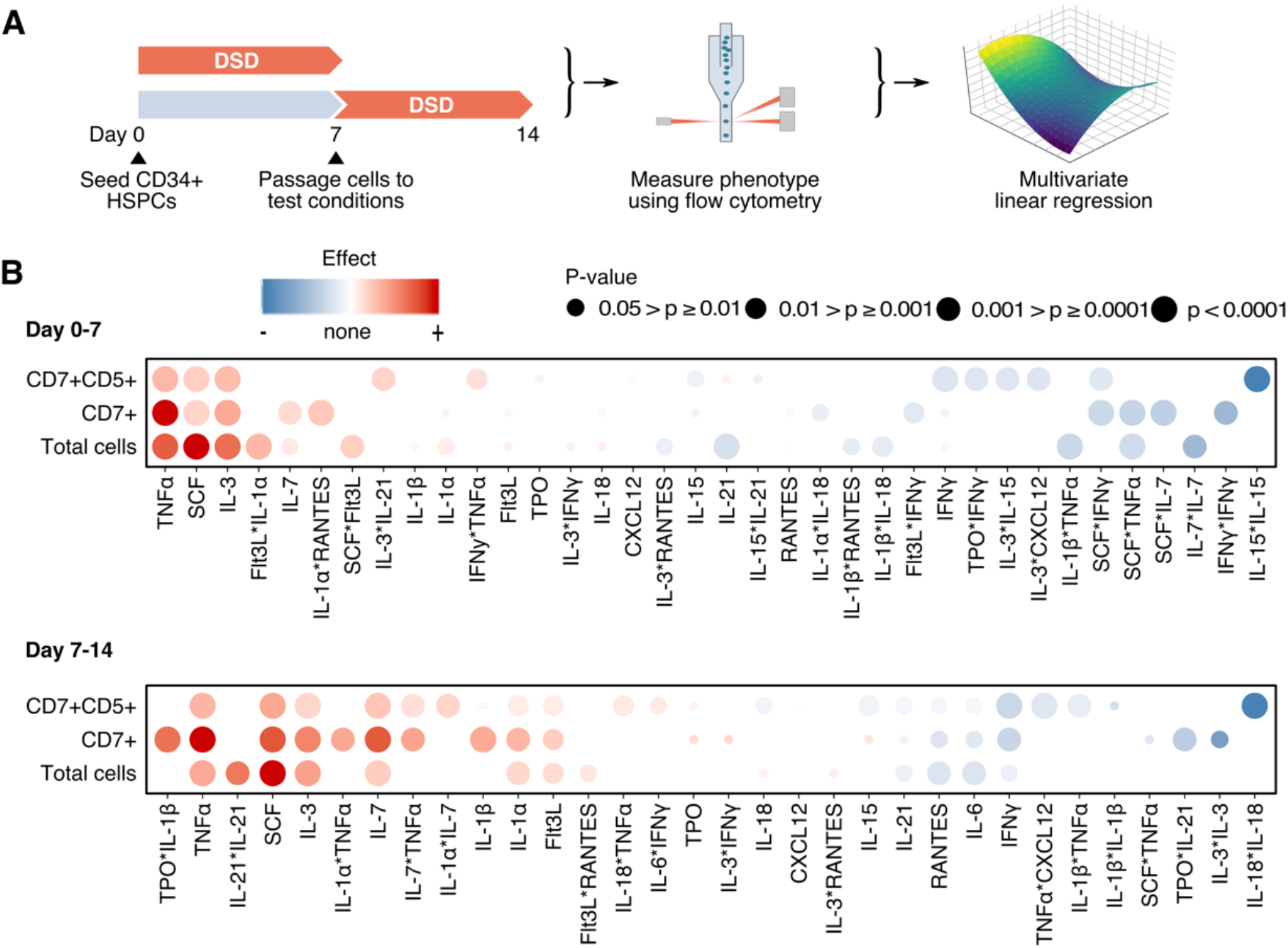
Screening for cytokines that enhance proT-cell differentiation and expansion. (A) Summary of two-part screening experiment workflow. Cells were cultured in screening conditions from day 0-7 or cultured until day 7 and passaged at equal densities into test conditions. These cells were then cultured until day 14. Cells from day 7 and 14 were harvested and analyzed using flow cytometry. The absolute count of each population of interest was measured and used to calculate a z-score relative the the control. The z-scores were then used to fit multivariate linear regression models. (B) Effect of cytokines on total cells, CD7^+^ lymphocytes, and CD7^+^CD5^+^ cells. Red indicates an effect greater than the control condition while blue indicates an effect lesser than the control. The size of the circle indicates the significance of the effect in the regression model. From n=2 independent UCB donors.

**Figure 2.**
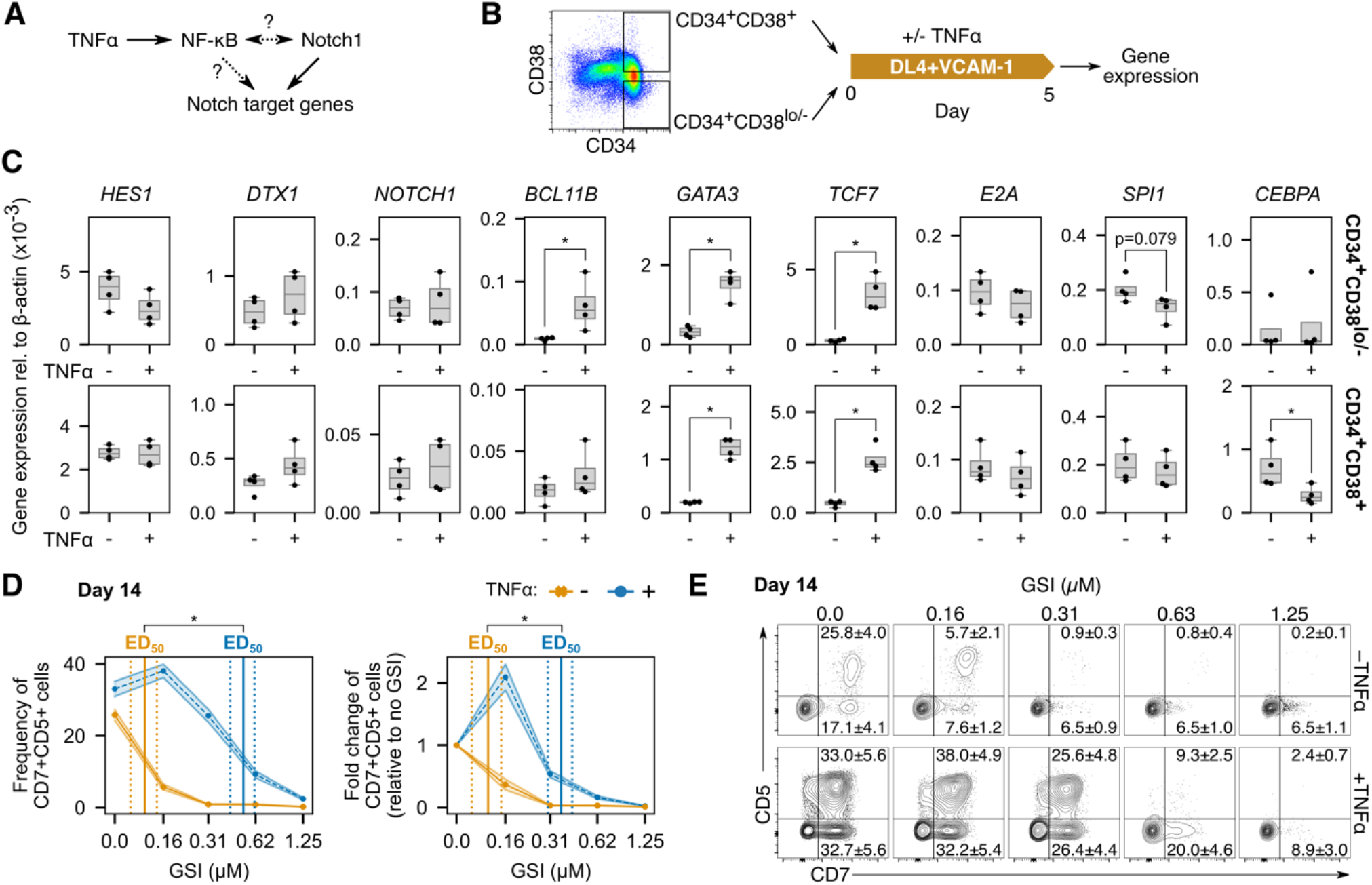
Synergy between TNFα and the Notch pathway compensates for low Notch activation. (A) TNFα activates the NF-κB pathway, which may regulate Notch target genes or regulate Notch itself. (B) To investigate the ways that TNFα may be interacting with Notch, CD34^+^ HSPCs were sorted into CD38^lo/-^ and CD38^+^ fractions and seeded separately on DL4+VCAM-1 with and without TNFα. Gene expression was measured using qPCR after 5 days of culture. (C) Both the CD38^lo/-^ and CD38^+^ fractions upregulated *GATA3* and *TCF7* in response to TNFα while only CD38^lo/-^ HSPCs upregulated *BCL11b*. No differences were observed in any other Notch target genes, implying that TNFα is not regulating Notch itself. CD38^lo/-^ but not CD38^+^ HSPCs downregulated *SPI1* slightly in response to TNFα. In contrast, only the CD38^+^ fraction significantly downregulated *CEBPA* when cultured with TNFα. From n=4 independent UCB donors. (D) CD34^+^ HSPCs were seeded on DL4+VCAM-1 for 14 days with increasing concentrations of γ-secretase inhibitor (GSI) to inhibit Notch activation. TNFα was able to maintain CD7^+^CD5^+^ cell generation with a signifantly higher concentration of GSI than the without. (E) Representative flow cytometry plots show the differential effects of Notch inhibition with and without TNFα. Shown are mean±standard error from n=7 independent UCB donors and *p<0.05.

### Flow Cytometry Analysis of Surface Markers

Adherent cells were collected from DL4+VCAM-1 surfaces using vigorous pipetting or enzymatically dissociated using TrypLE Express (ThermoFisher). Samples from Figures 1–3 were stained and collected with a LSRFortessa cytometer (BD) as previously described^20^. Compensation and gating was with FlowJo X software. The antibodies used in these experiments are listed in Supplemental Table 38. Samples in Figure 4 were collected with a CytoFlex cytometer (Beckman Coulter). Collected cells were rinsed twice with PBS and stained with Zombie-UV viability dye (BioLegend). Cells were then stained with fluorophore-conjugated antibodies diluted in Brilliant Stain Buffer (BD). Finally, cells were rinsed to remove unbound antibody and resuspended in Hanks Balanced Salt Solution (ThermoFisher) with 2% FBS (Gibco) for analysis. Compensation was performed with the cytometer software (CytExpert v2.3) and gating was performed with FlowJo X. The antibodies used are listed in Supplemental Table 39. Further analysis and statistics was with the R (version 3.3.2) or Python (version 3.7) programming languages.

**Figure 3.**
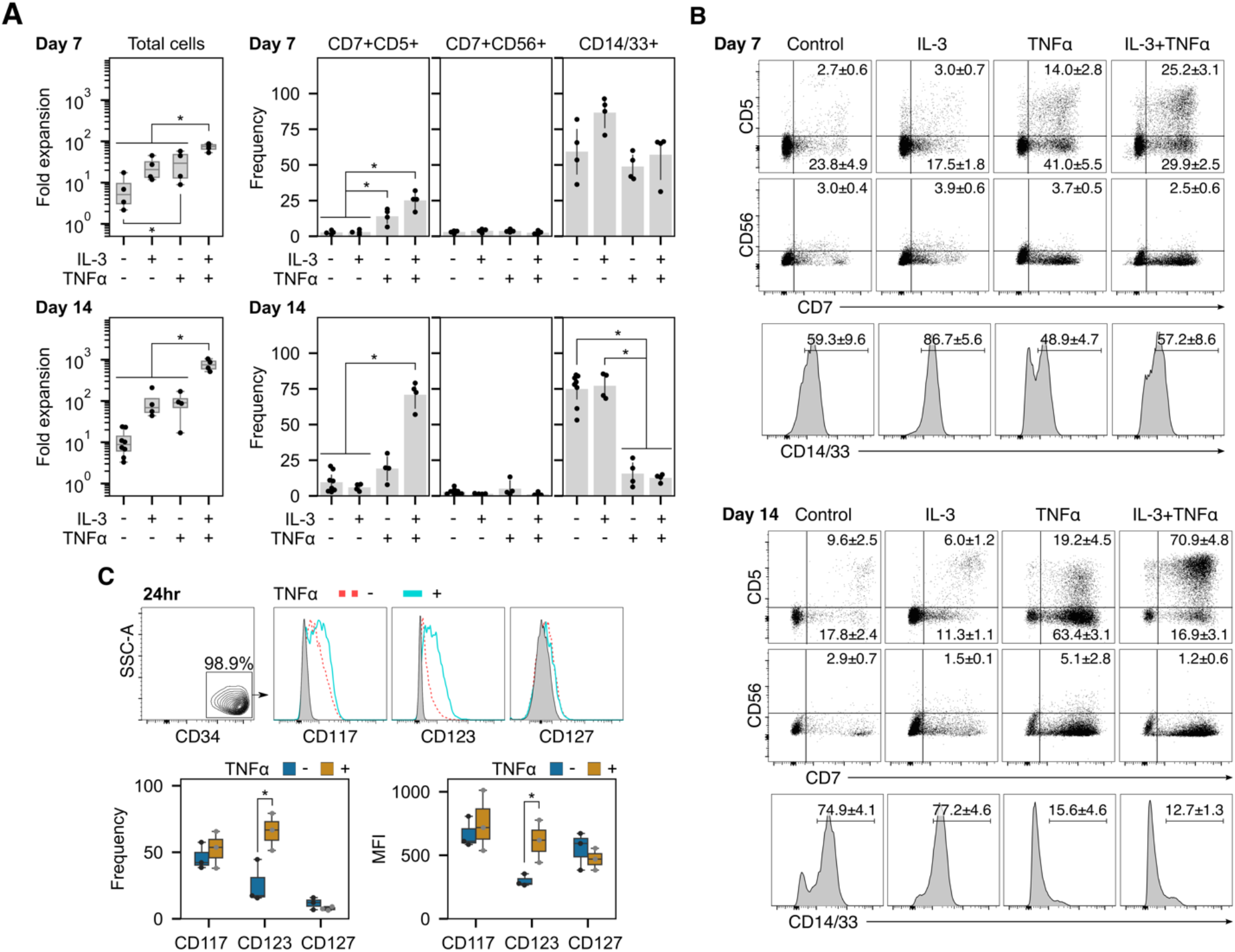
IL-3 and TNFα enhance proT-cell expansion and purity. CD34^+^ HSPCs were placed on DL4+VCAM-1 and fold expansion and phenotype measured on day 7 and 14. (A) Fold expansion and frequency of CD7^+^CD5^+^ proT, CD7^+^CD56^+^ NK, and CD14/33^+^ myeloid cells on day 7 and 14. Combining TNFα with IL-3 significantly increased total cell expansion over all other conditions. It also increased the frequency of CD7^+^CD5^+^ cells without increasing CD7^+^CD56^+^ frequencies. By day 14, the frequency of CD14/33^+^ cells was also significantly lower in groups containing TNFα than those without. (B) Representative flow cytometry plots from day 7 and 14 show a relatively pure population of CD7^+^CD5^+^ cells when TNFα is combined with IL-3. Frequencies are mean ± standard error from n = 4 independent UCB donors and *p<0.05. (C) CD117, CD123, and CD127 expression on CD34^+^ HSPCs with or without TNFα stimulation for 24 hours. TNFα induced a significant increase in the frequency of CD123^+^ cells. The increased frequency of CD123^+^ cells was accompanied by an increase in the median fluorescent intensity (MFI) of CD123, indicating a higher receptor density on cell’s surface after TNFα stimulation. *p<0.05 for n=3 independent UCB donors.

**Figure 4.**
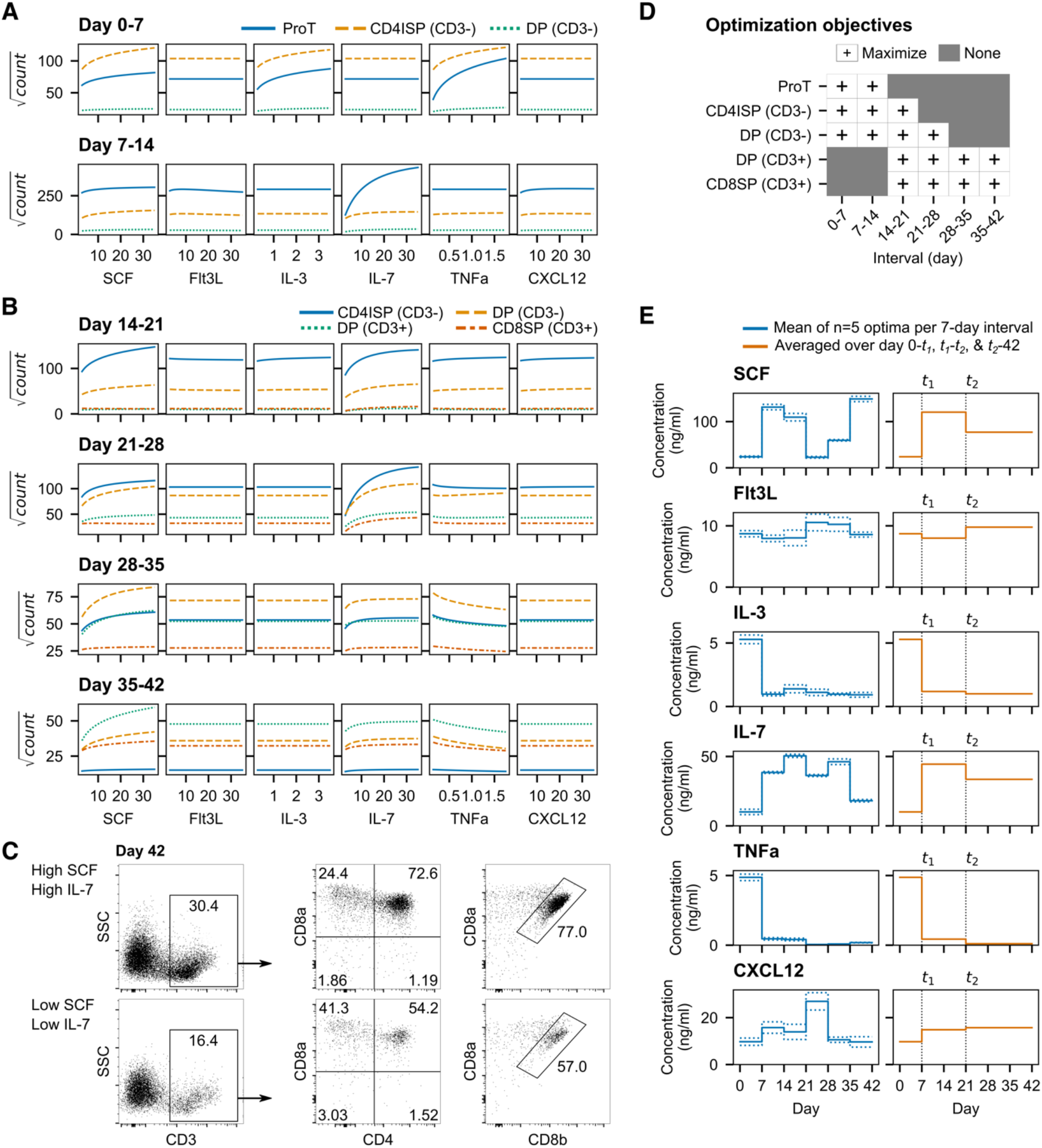
SCF and IL-7 enhance T-cell maturation while TNFα inhibits maturation. (A) Dose response of proT, CD4ISP, and early DP to test cytokines between day 0-7 and 7-14. Cells are unresponsive to IL-7 until day 7-14 but respond strongly to SCF, IL-3, and TNFα from day 0-7. (B) Dose response of CD4ISP, early and late DP, and CD8SP to test cytokines for each 7-day interval between day 14-42. The positive dose-dependent effect of IL-3 and TNFα early in cultures flattens and TNFα begins to inhibit the generation of DP and CD8SP cells. In (A-B), cytokine concentrations were swept from low to high while holding all other cytokines at their scaled center value (0). Shown are square root transformed cell counts for each population. RSM was constructed using n=3 pooled UCB donors. (C) Representative flow cytometry plots for day 42 showing sample conditions with either high SCF and IL-7 or low SCF and IL-7. High SCF and IL-7 increased the proportion of CD3^+^ T-cells, the majority of which were CD4^+^CD8ab^+^ or CD4^−^CD8ab^+^. (D) Objectives for optimization of RSM. Populations were either maximized or not included/present in certain 7-day intervals. (E) Optimized cytokines per 7-day interval (left) or as a three-stage assay (right). Solid lines are the mean of the top 5 optimizations while dotted lines represent the standard deviation.

### Proliferation Assays

Cells were stained with 2.5μM CellTrace CFSE (ThermoFisher) in PBS and incubated at 37°C for 8 minutes. CFSE was quenched by adding 5 times the volume of IMDM+BIT and incubating at 37°C for 5 minutes. Cells were then resuspended in fresh media and seeded on DL4+VCAM-1 for culture. For analysis, cells were collected and analyzed using a LSRFortessa cytometer. Proliferation was modelled using FlowJo X software and further analysis was performed with R.

### Quantitative PCR

Cells were lysed and RNA isolated using the PureLink RNA Micro Kit (Invitrogen) according to the manufacturer’s protocol. cDNA was reverse-transcribed from RNA using SuperScript III Reverse Transcriptase (Invitrogen) according to the manufacturer’s protocol. cDNA was amplified with primers and FastStart SYBR Green Mastermix (Roche) using QuantStudio 6 Flex (Applied Biosystems). Relative expression of individual genes was calculated by the delta cycle threshold (ΔCt) method and normalized to β-actin. PCR primer sequences are available in Supplemental Table 40.

### Cytokine Screens

For screening experiments, a Definitive Screening Design (DSD)^21^ was employed where each cytokine was tested at three concentrations. Cytokine concentrations for initial screening experiments are shown in Supplemental Table 1 and for follow-up experiments in Supplemental Table 8. Absolute cell numbers were acquired for each cell population using flow cytometry and, for initial experiments, a z-score was calculated relative to a control condition in order to gain a better- or worse-than estimate. The control condition was the cytokines used previously in the DL4+VCAM-1 assay as in Shukla *et al*.^20^. For follow up experiments, the regression model was fit to the cell counts instead of z-scores. Additional information about screening experimental design and analysis can be found in Supplemental Methods.

### Response Surface Methodology (RSM) Models of Cytokine Dose Responses

RSM experiments were orthogonal central composite designs (CCD) comprising 6 cytokines at 5 concentrations (Supplemental Table 15). Throughout the experiment cells were passaged every 7 days onto freshly DL4+VCAM-1 coated plates in 100μl of fresh media. An additional 100μl of media was added in between passages. Absolute cell counts were acquired for all populations of interest using flow cytometry and multivariate linear regression was used to construct the RSM models. Additional information about RSM experimental design and analysis can be found in Supplemental Methods.

### Optimizing RSM Models for T-cell Maturation

Regression coefficient estimates were used to build models in custom Python code (version 3.7). Desirability functions were used to maximize multiple RSM models simultaneously using the Basin-Hopping algorithm in the *SciPy* library^22,23^. Additional information about the optimization procedure can be found in Supplemental Methods.

### Statistical Analysis

With the exception of screening and RSM experiments, all statists were calculated in R (version 3.3.2). A Shapiro-Wilks normality test was used to determine whether data could be appropriately modeled by a Gaussian distribution. If data was non-Gaussian (Shapiro-Wilks p<0.05), a non-parametric Kruskal-Wallis test with Dunn’s post-hoc analysis was used, and the false discovery rate was minimized using the Benjamini-Hochberg p-value adjustment. Otherwise, one-way ANOVA with Tukey post-hoc analysis was employed.

### Data Sharing Statement

Regression coefficient estimates for DSD and RSM experiments are provided in a data supplement available with the online version of this article. The custom Python code used to build and optimize RSM models from coefficient estimates is available at gitlab.com/stemcellbioengineering/polynomialfeatures. For original data, please contact.

## RESULTS

### A Two-Phase Screening Strategy for Enhancers of ProT-Cell Differentiation

Using the DL4+VCAM-1 platform, we sought to identify soluble cytokines that positively regulate T-lineage differentiation and expansion from HSPCs. A list of 15 candidate molecules was assembled from the literature and screened for total, CD7^+^ lymphocyte, and CD7^+^CD5^+^ proT-cell expansion. To separate the effects of cytokines on early HSPCs and emerging proT-cells, the experiment was performed in two separate stages (Figure 1A). Test conditions were compared to a control condition was included that contained SCF, Flt3L, TPO, and IL-7 (4F) each at 100 ng/ml, as was used in our previous work^20^.

Of the 15 cytokines tested, SCF, IL-3, and TNFα elicited strong proliferative effects from day 0-7 on all three populations while IL-7 had only a small effect on total and CD7^+^ cell numbers (Figure 1B). From day 7-14, the effect of IL-7 was much greater, although cells still responded most strongly to SCF, IL-3, and TNFα. Other cytokines, such as IFNg and IL-6, had a negative effect on expansion of one or more of the cell populations, and were excluded from future experiments. A similar experiment was performed including IL-3 and TNFα at a higher range of concentrations in order to confirm our observations (Supplemental Figure 2). From this, working concentrations for IL-3 and TNFα were chosen as 10 and 5 ng/ml, respectively.

We measured proliferation using carboxyfluorescein succinimidyl ester (CFSE) dye for 4F cytokines (control) or 4F plus one of IL-3 and TNFα. Cells treated with IL-3 proliferated more than the control but this was primarily in the non-lymphoid (CD7^−^) fraction while cells treated with TNFα proliferated similarly to the control (Supplemental Figure 3A-B). All cells transitioned through proT1 and proT2 stages after 5 days, consistent with development on OP9-DL4^5^, though a significantly (p<0.05) higher proportion of CD7^+^ cells treated with TNFα had a proT2 phenotype (37.6±4.9%) compared to the IL-3 (17.5±2.1%) and control (15.2±2.6%) (Supplemental Figure 3C).

### Interactions Between TNFα and the Notch Pathway Enhances T-Lineage Differentiation

The early increase in CD7^+^ expression with TNFα made us ask whether it was interacting with the Notch1 pathway to enhance T-lineage specification. TNFα signals through the NF-κB pathway which, in other cell types, has been shown to interact extensively with Notch in a context-dependent manner (Figure 2A)^24^. We were interested to see if the effects of TNFα were specific to HSCs and multipotent progenitors (MPPs) or their downstream progeny. We therefore sorted CD34^+^ HSPCs into CD38^lo/-^ and CD38^+^ fractions to separate HSCs/MPPs from more differentiated progenitors. We seeded each fraction on DL4+VCAM-1 with and without TNFα and measured the expression of Notch target genes after 5 days (Figure 2B). The addition of TNFα increased the expression of *GATA3* and *TCF7* (encoding TCF-1)—genes that are important for T-lineage specification^25^—in both the CD38^lo/-^ and CD38^+^ fractions (Figure 2C). Additionally, *BCL11B*, a gene important for T-lineage commitment, was upregulated in CD38^lo/-^ cells treated with TNFα. No significant differences were observed in *HES1, DTX1, E2A*, or *NOTCH1* mRNA levels, suggesting that this effect was not due to an increase in Notch1 receptor expression and overall Notch pathway activation. TNFα also induced a modest decrease in the myeloid gene *SPI1* (encoding PU.1) in CD38^lo/-^ HSPCs, and a significant decrease in *CEBPA* in CD38^+^ HSPCs. The decrease in *CEBPA* mRNA levels was not due to an increase in *HES1* expression, which antagonizes *CEBPA^26^*. The upregulation of T-lineage genes and decrease in pro-myeloid-lineage genes by TNFα provides a mechanism by which it inhibits myeloid differentiation. The increased expression of *BCL11B* in only the CD38^lo/-^ fraction implies that they have a higher propensity for T-lineage differentiation, consistent with previous reports^5^.

Given that TNFα enhances the expression of T-lineage specification genes, we next tested whether it could decrease dependence on Notch signaling during T-cell development. We placed CD34^+^ HSPCs on DL4+VCAM-1 for 14 days and used the γ-secretase inhibitor (GSI) DAPT to inhibit Notch activation. We calculated the 50% effective dose (ED_50_) of GSI for CD7^+^CD5^+^ cell with and without TNFα using linear interpolation. Without TNFα, the ED_50_ for the frequency of CD7^+^CD5^+^ cells was 0.12±0.02μM (mean±standard error) and with TNFα was 0.52±0.10μM (Figure 2D-E). Likewise, the ED_50_ for the fold-change in CD7^+^CD5^+^ cell numbers without TNFα was 0.12±0.03μM and with TNFα was 0.36±0.06μM. Thus, TNFα can partially compensate for Notch pathway inhibition through the regulation of Notch target genes although it cannot replace Notch activity completely.

### TNFα Synergizes with IL-3 to Enhance ProT-cell Expansion

On its own, IL-3 had little effect on overall CD7^+^CD5^+^ cell expansion, and preferentially expanded non-lymphoid (CD7^−^) cells early in cultures. Given its positive effect on CD7^+^CD5^+^ cell development in screening experiments, we hypothesized that IL-3 might interact synergistically with TNFα. We therefore seeded CD34^+^ HSPCs on DL4+VCAM-1 with IL-3+TNFα in combination. The use of both cytokines led to significantly higher expansion than with any single cytokine alone, and cells were confluent by day 7 and required passaging.

By day 7, the IL-3+TNFα group had expanded 75.0 (55.6-86.8)-fold (median, 5-95 percentile) compared to 20.7 (12.2-42.7)-fold in IL-3, 29.5 (9.74-55.9)-fold in TNFα, and 5.1 (2.4-15.9)- fold in the control, and had a higher frequency of CD7^+^CD5^+^ cells than IL-3 and control (Figure 3A). CD7^+^CD56^+^ NK frequencies were less than 4% in all groups and the frequencies of CD14/33^+^ myeloid cells were variable and not significantly different. By day 14, fold expansion was an order of magnitude greater in the IL-3+TNFα group at 753.2 (532.4-1026.9)-fold compared to 69.0 (45.8-190.8)-fold in the IL-3, 90.7 (27.5-159.1)-fold in the TNFα, and 8.9

(4.3-21.5)-fold in the control. The frequency of CD7^+^CD5^+^ cells was also significantly greater with IL-3+TNFα than all other conditions. As on day 7, CD7^+^CD56^+^ NK frequencies were low and both TNFα and IL-3+TNFα groups had a significantly lower frequency of CD14/33^+^ myeloid than IL-3 or control groups. Thus, IL-3+TNFα synergize to elicit significant proliferation and preferentially enrich cultures for CD7^+^CD5^+^ cells.

### TNFα Regulates Expression of the IL-3 Receptor

To elucidate a mechanism for the synergistic effect of IL-3 and TNFα, we examined the expression of the IL-3 receptor (CD123) after stimulation of CD34^+^ HSPCs with TNFα. TNFα has been reported to upregulate CD123 in human bone marrow (BM)-derived CD34^+^ HSPCs^27^. Consistent with this, we found that TNFα increased the frequency of CD123^+^ cells after 24h (Figure 3C). In addition, TNFα stimulation also increased the median fluorescent intensity (MFI) of CD123 expression, indicating an increase in the number of CD123 molecules on cell’s surface. We observed no change in the frequency and MFI of the SCF (CD117) and IL-7 (CD127) receptors in our UCB-derived HSPCs. Thus, the synergy between IL-3 and TNFα is due, at least partly, to increased responsiveness of cells to IL-3 through an increase in the number of cells expressing the receptor and well as an increase in the level of expression.

### TNFα switches from an enhancer to an inhibitor during T-cell development

Next, we sought identify cytokine signaling requirements for T-lineage maturation on DL4+VCAM-1. We used response surface methodology (RSM) to model the dose response of cells to SCF, Flt3L. IL-3, IL-7, TNFα, and CXCL12 (Supplemental Figure 4A). Regression was used to fit polynomial models to experimental data (Supplemental Figure 5). We excluded TPO after we found that removing it reduced CD14/33^+^ myeloid generation without detrimentally affecting CD7^+^ lymphoid expansion (Supplemental Figure 6). CXCL12 was included because we observed a modest but positive effect in screening experiments and its reported positive effect on cell survival during β-selection^28,29^. Following our Notch inhibition experiment and reports that aβT-cell development requires reduced Notch pathway activation around the β-selection checkpoint^30^, we titrated DL4 and reduced the concentration 7.5-fold while maintaining similar proportions of proT-cells (Supplemental Figure 7). Experiments were conducted over 7-day intervals and the number of cells in each population measured using flow cytometry. Because the RSM has higher predictive power than the definitive screening design, we included the first two weeks of differentiation to more accurately estimate cytokine dose responses during T-cell specification. We measured proT-cells, CD4ISPs, and early DPs (CD3^−^) during the first 14 days. From day 14 onwards we included CD4ISPs, early DPs, late DPs (CD3^+^), and CD8SPs (CD4^−^ CD8a^+^CD3^+^).

On day 7 a large populations of proT and CD4ISP were present in cultures. These cells responded strongly to increasing concentrations of SCF, IL-3, and TNFα. Positive two factor interactive effects were observed between SCF and TNFα and IL-3 and TNFα (Supplemental Figure 8A). A small number of early DP cells were also present by day 7. Between day 7-14, all populations were responsive to increasing concentrations of SCF and IL-7 and unresponsive to IL-3. Cells responded positively to TNFα and CXCL12 with a negative interaction but the magnitude of these effects were small compared to SCF and IL-7 (Supplemental Figure 8B).

Cultures were primarily CD3^−^ on day 21 but the proportions of CD3^+^ cells steadily increased from day 28 onwards (Figure 4B). Again, TNFα had a small positive effect on cell expansion at higher concentrations between day 14-21 but this became negative from day 21 onwards. Differentiation was dominated by SCF and IL-7 and the cytokines had some interactive effects from day 21 through day 35 (Figure 4C and Supplemental Figure 8C-E). The response to IL-3 was small between day 21-28 and cells were completely unresponsive to IL-3 from day 28 onwards. Flt3L had little or no effect on cell expansion throughout the entire assay.

### An optimized three-stage assay for *in vitro* T-cell generation

In order to define a set of preferred conditions for each step in the differentiation, we optimized the RSMs to find the cytokine concentrations that maximized the number of cells in different populations for each 7-day interval. A desirability function for each population was used to calculate an overall desirability which was maximized using the basin-hopping algorithm (Supplemental Figure 4B)^23^. The optimization objectives were changed throughout the differentiation to reflect the populations present, first maximizing cells in the CD3^−^ populations and shifting to CD3^+^ cells (Figure 4D).

The top five solutions from the optimization converged to the same overall desirability score indicating that a they are global maxima (Figure 4E and Supplemental Figure 4C). Next, we constructed a three-stage protocol that approximated the 7-day interval optima as closely as possible. We split the assay into the intervals [0, *t_1_*], [*t_1_, t_2_*,], and [*t_2_*, 42] days, where *t_1_, t_2_* are multiples of 7 in [7, 42) and *t_1_ < t_2_*. We used a brute force search method that averaged the predicted optima within the intervals for every *t_1_, t_2_* and found the pair that had the highest average overall desirability across time. Using this approach, we found *t_1_*=7 and *t_2_*=21 maintained desirability scores that were close to the 7-day interval scores (Supplemental Figure 4D). The updated three-stage cytokine concentrations are provided in Supplemental Table 37.

## DISCUSSION

Current clinically relevant protocols for generating T-cells from stem cells have, thus far, not been able to generate cells in the quantities or at the purity necessary to make them suitable for translation. Here, we show the benefit of using a defined system for isolating and studying the effects of cytokines on T-cell development and identified interactions between TNFα and the Notch pathway that enhance T-lineage specification. In addition, we found that combining TNFα with IL-3 strongly potentiated proT-cell proliferation, independent of effects on myeloid lineage cells (Figure 5A). We then optimized cytokine concentrations for maturation to generate CD3^+^ DP and CD8SP T-cells *in vitro* and provide an updated DL4+VCAM-1 assay as a three-stage protocol.

**Figure 5.**
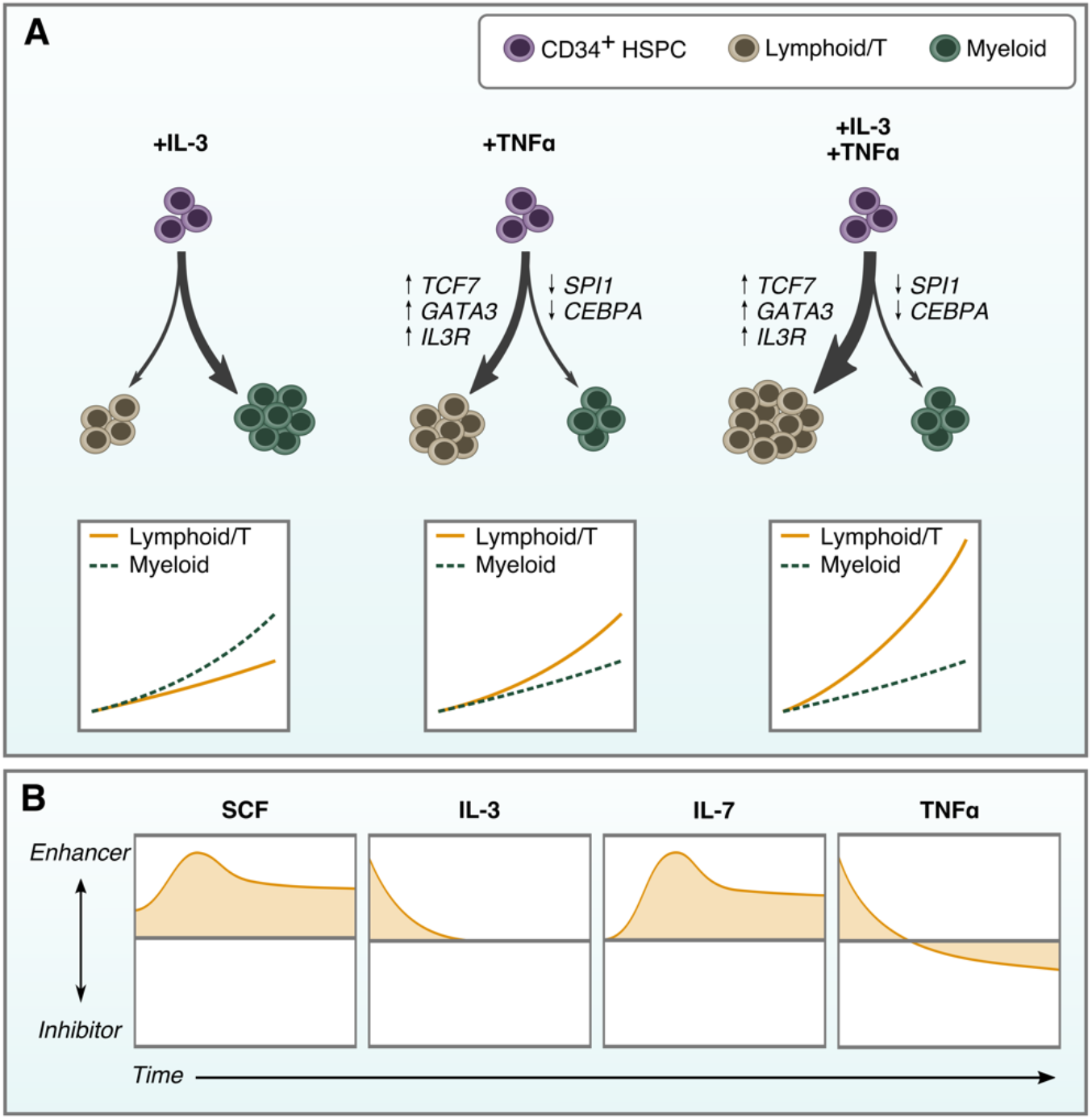
TNFα couples T-lineage differentiation with IL-3-induced proliferation. (A) Individually, IL-3 stimulates proliferation of myeloid-biased CD34^+^ HSPCs while TNFα induces T-lineage differentiation by positively regulating Notch target genes. It also increases the proportion of cells responsive to IL-3 by regulating the IL-3 receptor. When combined, IL-3 provides strong proliferative signals to developing lymphocytes, leading to substantial expansion of T-lineage cells. (B) The positive effects of IL-3 and TNFα are on T-cell specification where cells have limited responsiveness to IL-7. Once T-lineage development is initiated, SCF and IL-7 become the drivers of proliferation, and responsiveness to IL-3 and TNFα is lost. TNFα becomes inhibitory when cells enter developmental stages associated with positive and negative selection

TNFα was previously reported to accelerate the differentiation of DN1 to the DN2 stage in murine thymocytes but the mechanism for this effect was not investigated^12^. Our findings show that TNFα has a similar effect on human T-lineage development through the upregulation of Notch target genes *GATA3, TCF7*, and *BCL11b*, and the downregulation of *SPI1* and *CEBPA*. The inhibitory effect of TNFα in later stages of T-cell development is likely due to increased apoptosis in cells undergoing selection^31,32^. TNFα is constitutively expressed in the human postnatal thymus, primarily in the cortico-medullary junction (CMJ) and medulla^33^. Thus, we expect progenitors entering the thymus near the CMJ and late DP/SP cells undergoing negative selection in the medulla to experience higher levels of TNFα than those in the cortex. Our findings that TNFα has a positive effect on cells during T-cell specification but a negative effect on later stages of development is consistent with thymus expression.

The combination of IL-3 and TNFα provided a significant enhancement in cell expansion and development to the T-lineage (Figure 5B). We assume that the signals provided by IL-3 and TNFα are orthogonal – that they regulate non-overlapping gene expression that results in a multiplicative increases in proliferation rather than simply additive. The increase in CD123 expression by TNFα likely mediates the proliferative effect, allowing for increased activation by IL-3 of the Janus kinase/signal transducer of and activator of transcription (JAK/STAT) and PI3K/Akt signaling pathways that regulate cell survival and proliferation^34–36^. IL-7 also signals via JAK/STAT and PI3K/Akt and there is likely overlap in IL-3 and IL-7 induced gene transcription^34^. On its own, IL-3 promoted the early expansion of myeloid lineage cells and TNFα mitigated this effect, likely through its interactions with the Notch pathway. IL-3 is produced by human TECs^37^ and, when combined with SCF, IL-1, IL-6, and IL-7, enables enhanced reconstitution of fetal thymic organ cultures (FTOCs) by mouse proT-cells^10^. Thus, it likely plays a physiological role in human T-lineage development in the thymus.

SCF’s role in proT-cell proliferation and differentiation is well-attributed but, to our knowledge, its positive effect on the later DP stages is unreported. SCF was shown to promote murine DN1-3 proliferation while inhibiting differentiation to the DP stage in OP9-DL1 co-cultures^38^. This may be due to differences between human and mouse T-cell development or simply an artifact of the assays in which they were studied (OP9-DL1 versus our defined system). Flt3L had a modest effect on T-cell development. It has been reported to promote survival of bone marrow-resident thymic precursors and a modest increase in murine proT-cell expansion in FTOCs, but our results suggest it has little effect on T-cell development in this system^39,40^.

Our work demonstrates the usefulness of building quantitative models using a fully defined culture system. As temporal changes in responses to cytokines and growth factors are challenging to study, we split experiments into discrete intervals that could be tested individually and then combined to approximate temporal signaling regimes. This required some prior knowledge to estimate the length of intervals necessary to capture the dynamics in question.

Shorter intervals would give finer resolution but at increased experimental workload. They may also miss developmental processes that occur over longer times. In the future, our RSM models could be augmented with data from shorter or longer experiments at different stages of development in order to capture fast developmental transitions and long-term cytokine dependencies.

Future work should explore *in vitro* T-cell maturation in cells that have been genetically engineered to express a CAR or antigen-specific TCR. We expect this would block endogenous TCR recombination through allelic exclusion while providing a free pass on positive selection that avoids contraction of cell numbers due to recombination of non-functional TCRs^41,42^. This would further accelerate differentiation kinetics while reducing the risk of graft-versus-host disease to enable robust and clinically translatable T-cell immunotherapies from stem cells.

## ACKNOWLEDGEMENTS

Recombinant human DL4-Fc was kindly provided by Ashton Trotman-Grant and Juan Carlos Zúñiga-Pflücker. Fluorescence-activated cell sorting (FACS) was performed by the SickKids-UHN Flow and Mass Cytometry Facility. This study was supported through a Canadian Institute for Health Research (CIHR) Foundation Grant to P.W.Z. J.M.E. was supported by the Natural Sciences and Engineering Research Council of Canada (NSERC) Canadian Graduate Scholarship – Masters and Doctoral, the Ontario Graduate Scholarship, the University of Toronto Mary H. Beatty Fellowship, and the Zymeworks – Michael Smith Laboratories Fellowship in Advanced Protein Therapeutics. P.W.Z. is the Canada Research Chair in Stem Cell Bioengineering.

## AUTHOR CONTRIBUTIONS

J.M.E. and P.W.Z. conceptualized the research and wrote the manuscript. J.M.E. performed experiments.

## CONFLICTS OF INTEREST

Intellectual property related to this work is being evaluated under an option to license. P.W.Z. is a scientific founder and consultant of Notch Therapeutics, a biotechnology company developing stem cell-derived T-cell immunotherapies.

