## Supplemental Methods and Data for "Inflammatory Cytokines Regulate T-Cell Development from Blood Progenitor Cells in a Stage and Dose-Specifc Manner"

#### Definitive Screening Design (DSD) Experiments

DSD are a method to screen a large number of factors in a relatively small experiment size[1]. They are three-level designs of at least $2k + 1$ runs, where $k$ is the number of factors being tested and levels are coded as -1,0,1. The design comprises $k$ fold-over pairs, where one factor is set at the mid-range (0) and all other factors are set to the extrema (-1 or 1). An added centre run with factors set to 0 enables DSDs to estimate all quadratic effects $X_{ii}^2$. In a DSD, all main effects $X_i$ are independent of any two-factor interactions $X_i X_j$ and no two-factor interactions are confounded with each other, although they may be correlated. For designs of 6 or more factors, it is possible to fit a full quadratic model to any 3 factors. DSDs may also include orthogonal blocks, allowing variance due to biological replicates to not confound factor estimates[2]. Experimental data is fitted with a quadratic model to estimate responses to screened factors.

We designed a 15-factor DSD with blocking to screen for cytokines that induce proliferation during T-cell specification of CD34$^+$ HSPCs on DL4+VCAM-1 (Supplemental Table 1). Two sequential DSD experiments were performed. The first, from day 0-7 and the second from day 7-14. In the latter, cells were cultured in the mid-range (0) concentrations for the first 7 days then passaged at equal densities to the test conditions. The motivation for this was to separate effects on early hematopoietic progenitor cells from proT-cells. Thus, it would allow us to identify if a factor had a positive effect in one stage and a negative effect in the other. As a control, we included a condition using the cytokines SCF, Flt3L, TPO, and IL-7 at the concentrations used in Shukla *et al*[3]. To get a better- or worse-than estimate of cytokine effects relative to the control, a z-score was calculated for each test condition:

$$z = \frac{test - control}{SE(control)}$$

where $SE(\cdot)$ is the standard error. At the end of each experiment, the number of total cells, CD7$^+$ lymphoid cells, and CD7$^+$CD5$^+$ proT-cells was measured using flow cytometry. Absolute cell counts were square root or log transformed as necessary to satisfy model assumptions that the residuals be approximately normally distributed. Due to the small size of DSD experiments, they rely on only a few factors being active. To avoid user bias in determining which factors to include, stepwise regression was used to automatically select the best model for the data. The minimum Bayesian information criterion (BIC) was used as the stopping rule during the procedure. BIC seeks to minimize the number of model parameters while maximizing the likelihood function, thus selecting the smallest model with the best fit possible. The value of the $\beta$ coefficient estimates for the resulting models were ranked by magnitude and visualized in order to determine which cytokines elicited the strongest effects on the populations measured. All parameter estimates are provided in Supplemental Tables 2-7.

We also constructed a 10-factor DSD as a follow-up experiment to confirm our results and to estimate working concentrations for IL-3 and TNFα. The same procedure was applied except that the relative cell counts were used instead of the z-score. The cytokines used and their concentrations are provided in Supplemental Table 8 while the parameter estimates are provided in Supplemental Tables 9-14.

**Response Surface Methodology (RSM) Models of Cytokine Dose Responses**

Our objective in using RSM was to find a quadratic approximation of the non-linear dose response of developing T-cells to cytokines. Recognizing that responsiveness to cytokines may change as cells differentiate, we designed successive 7 day experiments where cells were cultured on DL4+VCAM-1 with different concentrations of test cytokines. At the end of 7 days we measured the absolute cell count for each population of interest using flow cytometry. We repeated this until day 42 for a total of six 7-day intervals. For each interval after day 0-7, a pool of cells were cultured on DL4+VCAM-1. When beginning an RSM experiment these cells were passaged at equal density to the test conditions in order to measure the relative growth expansion for each condition. Three populations were measured in the first 14 days: proT ($CD7^+CD5^+$), CD4ISP ($CD4^+CD8a^-CD3^-$), and early DP cells ($CD4^+CD8a^+CD3^-$). From day 14 onward we looked at early DP, late DP ($CD4^+CD8a^+CD3^+$) and CD8SP ($CD4^-CD8a^+CD3^+$) cells. A small number of CD4SP ($CD4^+CD8a^-CD3^+$) cells were present at later timepoints but were infrequent and were thus excluded. All populations were mutually exclusive except for proT-cells which could potentially overlap with any of the other populations. However, our experience is that most cells become $CD7^{low/-}$ as they start to express CD4. Therefore, the proT-cell population does not completely overlap with any of the others.

In order to fit a quadratic function to experimental data, a 5-level, 54 condition central composite design (CCD) for 6 factors with orthogonal blocking was constructed from a $2^{6-1}$ fractional factorial experiment (32 conditions) with added centre (10 conditions) and axial points (12 conditions). The concentrations of each cytokine were coded: (-1, 1) for the fractional portion, 0 for the centre points, and $\alpha$=2.366 for the axial points. Axial points are where one cytokine is set to $\pm\alpha$ while all other cytokines are set to 0. Combined with the centre points where all cytokines are set to 0, they enable the estimation of quadratic dose responses. Scaled levels could be converted to cytokine concentrations by $c = c_0 * 3.5^X$, where $X$ is the scaled level and $c_0$ is the concentration when $X = 0$. This ensured that concentrations were continuous so that interpolated levels could be converted to physical concentrations. The value of $c_0$ for each cytokine was set to ensure that the working concentrations used in previous experiments were within range (Supplemental Table 15). The exponent base (3.5) was chosen to ensure that the range of cytokines was wide enough so that the change in response would be greater than measurement noise or biological variation.

A quadratic model was fit to each population for each time interval that has the form:

$$\hat{Y} = \beta_0 + \sum_{i=1}^{k} \beta_i X_i + \sum_{i=1}^{k-1} \sum_{j=i+1}^{k} \beta_{ij} X_i X_j + \sum_{i=1}^{k} \beta_{ii} X_{ii}^2 + \epsilon$$

where $\hat{Y}$ is the response variable (number of cells), $X$ are the coded factor levels, and $\beta$, $\epsilon$ are the parameters to be estimated using regression. The $X_i$ terms are main effects, $X_i X_j$ are two-factor interactions where the effect is greater than additive, and $X_{ii}^2$ are quadratic terms that result in a non-linear response. Accurately predicting model parameters assumes that residuals be approximately normally distributed. To ensure this, we transformed the absolute cell counts using a square root transform $\hat{Y} = \sqrt{Y}$. After fitting, the residual distributions for the models were fit with a normal distribution and a Shapiro-Wilks Goodness-of-Fit test was applied to the fitted distribution. A test value of $p>0.05$ indicated that the residuals were approximately normal.

Least-squares regression was used to estimate model parameters. For each time interval a multiple response model was used that fit the same parameters to all responses (cell populations). A full model was fit and then reduced by removing terms that were not significant to p<0.01, unless the non-significant term was part of a higher order effect. Plots of actual versus predicted values were used to check for potential outliers. In some cases, an axial condition for SCF and IL-7 was a notable outlier. In these instances, a new model was built that included a third order term ($X^3$) for that factor which improved the overall fit (Supplemental Figure 4). While the model terms were the same for all cell populations within a given time interval, models from different time intervals could use different terms. All parameter estimates are provided in Supplemental Tables 16-36.

**Optimization of RSM Models per 7 Day Interval**

The utility of regression models created using RSM is their ability to interpolate between test conditions to predict optimal responses. We were interested in optimizing these models to predict the cytokine concentrations that are best suited to T-cell maturation in the DL4+VCAM-1 assay. We assumed an ancestor-progeny relationship (proT → CD4ISP → early DP → late DP → CD8SP) where increasing the number of ancestor cells at time $t_n$ increases the number of progeny at time $t_{n+1}$. Therefore, if we are interested in producing as many late DP or CD8SPs as possible, we first want to find conditions that increase proT, then CD4ISP, then early DP numbers. However, differentiation is not synchronous and any of these populations may be present in cultures at the same time. This requires a multi-objective optimization strategy to find cytokine concentrations that best suit the growth and development of multiple cell types at once.

We used desirability functions to optimize multiple objectives simultaneously[4]. For each RSM model, we defined a piecewise objective function to maximize the response $\hat{Y}_i$:

$$d_i(\hat{Y}_i) = \begin{cases} 0 & \hat{Y}_i < L_i \\ \left(\dfrac{\hat{Y}_i - L_i}{U_i - L_i}\right)^s & L_i \leq \hat{Y}_i \leq U_i \\ 1 & \hat{Y}_i > U_i \end{cases}$$

where $U_i$, $L_i$ are upper and lower limits, respectively, that set the boundaries of the objective. This function scales the response to [0,1] where responses $\hat{Y}_i < L_i$ are undesirable and $\hat{Y}_i > U_i$ are most desirable. We set $L_i = 0$ and $U_i = 2\hat{Y}_{i,max}$, where $\hat{Y}_{i,max}$ is the maximum value when sweeping one factor at a time from low to high while holding the others at 0. When the scaling factor $s = 1$ the function is linear for $L_i \leq \hat{Y}_i \leq U_i$.

The overall desirability combines all individual desirability functions using the geometric mean:

$$D = \left(\prod_{i=1}^{p} d_i\right)^{\frac{1}{p}}$$

for $i = 1,2, \ldots, p$ objectives. Notice that for any $d_i(\hat{Y}_i) = 0$ the overall desirability is 0. The utility of this approach is that by combining multiple objectives into one single objective we can use any single objective optimization algorithm. We used the Basin-Hopping algorithm from the *SciPy* library in Python 3.7 (`scipy.optimize.basinhopping`) which is well-suited to multivariable multimodal optimization problems[5]. Beginning from a random initialization $X_0$, the algorithm cycles through the following steps:

1. Perform a random perturbation in $X$.
2. Local minimization using the Nelder-Mead simplex method.
3. Accept or reject the new $X$ based on the minimized function value $(-D)$.

Because the algorithm is a stochastic global optimizer, there is no way to know whether the solution returned is the true global minimum. Therefore, this procedure was repeated from 25 random initializations and the best 5 solutions were retained. To ensure that solutions were within the concentration levels tested, we constrained the search space using the $L2$-norm of $X$. By setting $D = 0$ when $\|X\|_2 > 2.366$, all solutions outside of the hypersphere design space were undesirable and therefore excluded. This prevented extrapolation beyond the concentrations tested where prediction uncertainty increases.

Each time interval was optimized separately using the above procedure. This yielded an optimal set of cytokine concentrations for each interval. Provided the optimizer converged on the global minimum, the overall desirability for the best 5 solutions are equal. However, this does not mean that the predicted cytokine concentrations will be the same. This is to be expected from and is typical for multi-objective optimization. These predicted cytokine concentrations comprise samples from the Pareto front – optimum solutions where one individual objective cannot be improved further without detrimentally affecting another objective. This is useful information for us because objectives with a larger variance in predicted optima indicate cytokines whose concentration is not as crucial as one with a small variance. Therefore, this gives us a method to qualitatively assess the importance of different cytokines at each interval in the differentiation.

**Optimizing Averaged Cytokine Concentrations Over Time**

The predicted optimum cytokine concentrations for each interval are accurate for the RSM models used in the optimization (provided the global minimum is found). However, the RSM models are statistical and their fit may be impacted by technical noise or measurement error that may result in fluctuations in predictions that are not biologically significant. Additionally, changing cytokine concentrations every 7 days is cumbersome to implement in the laboratory. To address both of these issues, we split the assay into three stages and selected the cytokine concentrations to use by averaging the 7-day optima within each stage. Rather than split the assay into three 14 day intervals, we opted to find those which would provide cytokine concentrations that were as close to the 7-day optima as possible. To do this, we split the assay into the intervals [0, $t_1$], [$t_1$, $t_2$,], and [$t_2$, 42] days, where $t_1$, $t_2$ are multiples of 7 in [7,42) and $t_1 < t_2$. For every pair of $t_1$, $t_2$ we averaged the optimal cytokine concentrations between the intervals and then calculated the overall desirability score for each 7-day interval using the averaged concentrations. This allows us to change cytokines concentrations as often as every 7 days during periods of rapid differentiation (ie. T-lineage specification) while periods of slower differentiation may use the same cytokine concentrations for 14 days or longer. Averaging over multiple time intervals also provides a consensus of optima to smooth out fluctuations due to noise.

### SUPPLEMENTAL FIGURES

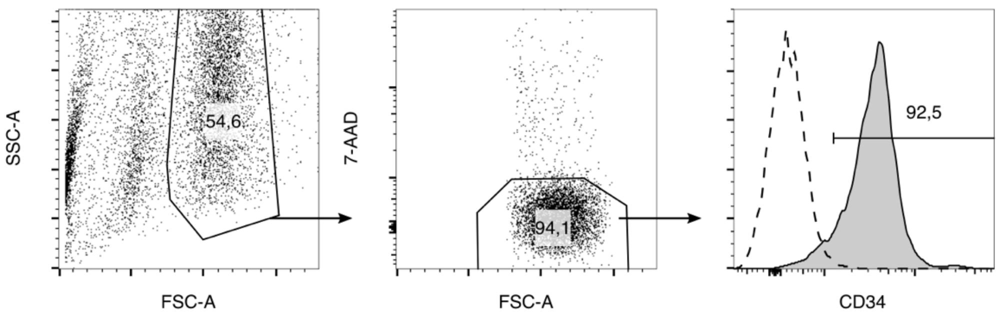


**Supplemental Figure 1.** Representative flow cytometry plot showing purity of umbilical cord blood-derived CD34+ HSPCs after isolation using EasySep magnetic beads.

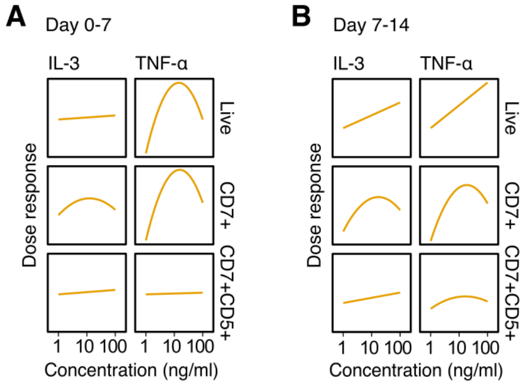


**Supplemental Figure 2.** Top cytokines from the initial screening experiment were used for second validation DSD experiment at a different range of concentrations. This was used to set the working concentration for IL-3 and TNFα to 10 and 5ng/ml, respectively. (A) IL-3 and TNFα dose response in cells from day 0-7. (B) IL-3 and TNFα dose response in cells from day 7-14. From n=2 independent UCB donors.

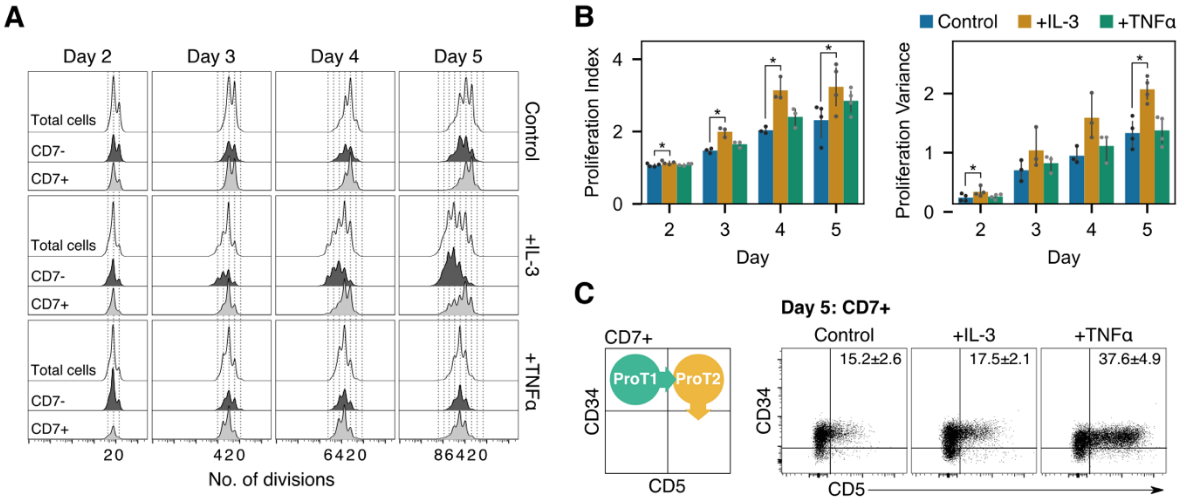


**Supplemental Figure 3.** Cell proliferation in response to IL-3 and TNFα. (A) Histograms of CFSE stained cells showing the number of divisions of each cell on day 2-5. Separation of $CD7^-$ and $CD7^+$ histograms shows differential responses to each cytokine. (B) Proliferation statistics from CFSE data. IL-3 treated cells had a larger proliferative index indicating that they underwent, on average, more divisions than the control group. They also had a significantly larger proliferative indicating that some cells responded much stronger to IL-3 than others. *p < 0.05 relative to the control on each day. (C) All groups transitioned through a proT1 to proT2 phenotype as expected during proT-cell development. Results are mean±standard error from n=4 independent UCB donors.

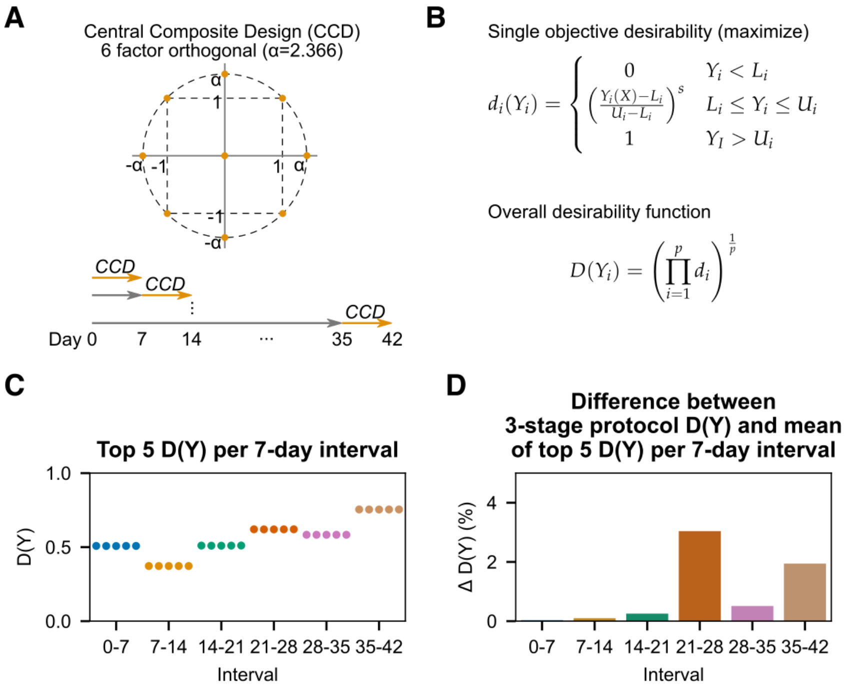


**Supplemental Figure 4.** (A) 2D representation of a central composite design and 7-day sequential RSM experiment design. (B) A single objective desirability function scales values of $Y_i$ between (0,1). The lower ($L_i$) and upper ($U_i$) set points provide a range for scaling and values that fall outside are set to 0 or 1, respectively. The overall desirability is calculated as the geometric mean of multiple single desirability functions. (C) The top 5 of 25 overall desirability scores were all equally desirable. (D) The overall desirability for each 7-day interval was calculated using the cytokine concentrations from the 3-stage design and compared to the mean of the top 5 desirability scores for each 7-day interval. The maximum difference was 3.0% for day 21-28 while most differences were <1%.

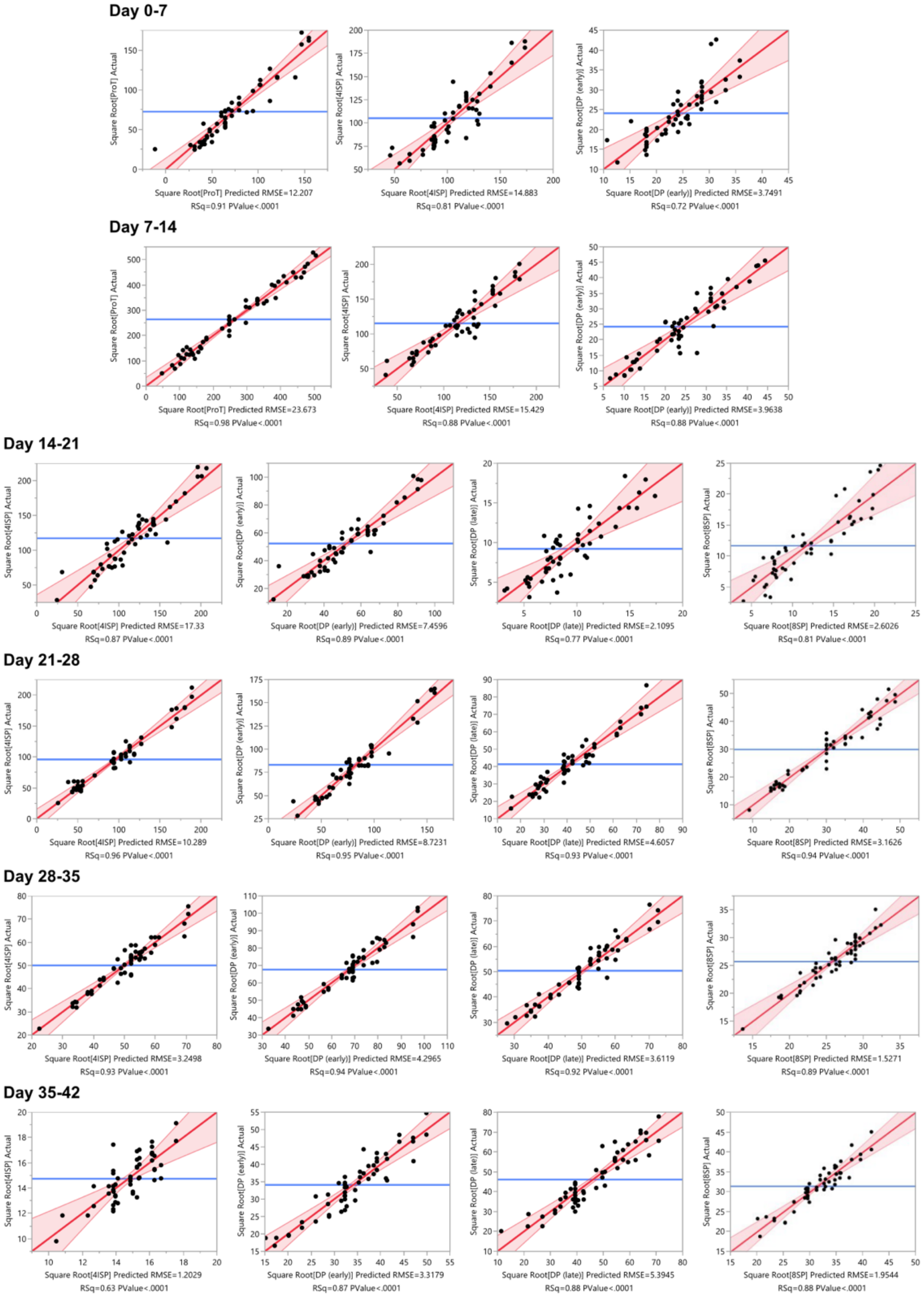


**Supplemental Figure 5.** Actual vs. predicted plots for RSM models. These were used to check for outliers and assess the quality of the model fit. DP (early) are $CD3^-$ and DP (late) are $CD3^+$.

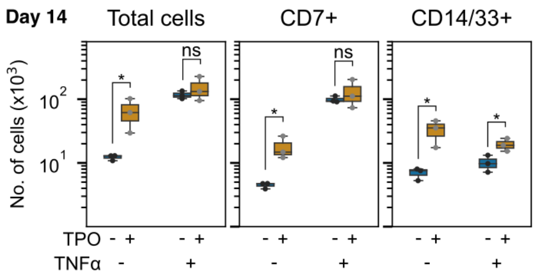


**Supplemental Figure 6.** Removing TPO from cultures containing TNFα reduces CD14/33$^{+}$ myeloid cell numbers without affecting total or CD7$^{+}$ cell numbers. *p<0.05 for n=3 independent UCB donors.

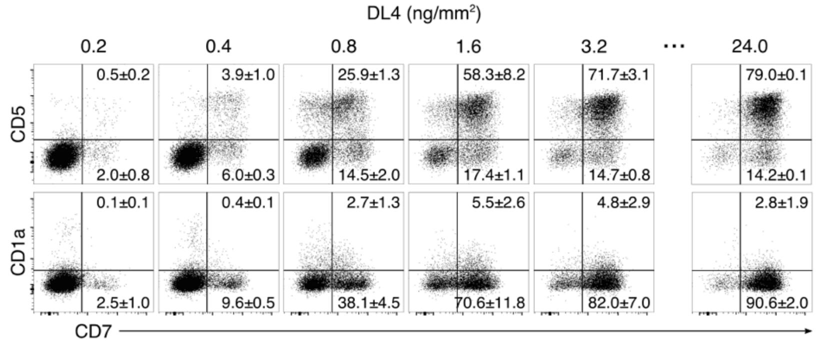


**Supplemental Figure 7.** IL-3+TNFα support T-lineage development with a 7.5-fold reduction in DL4 coating concentration. From n=2 independent UCB donors.

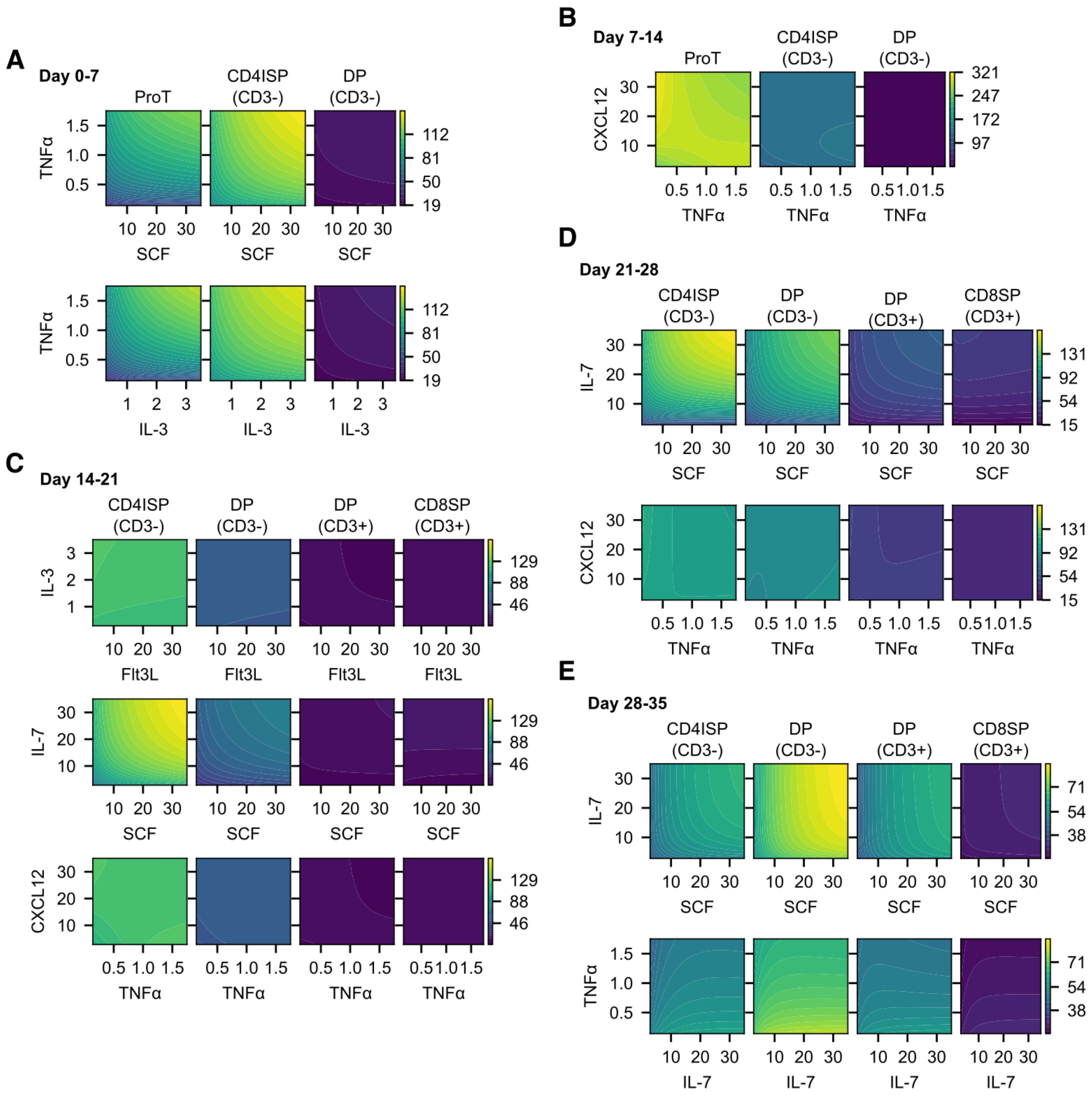


**Supplemental Figure 8**. Two factor interactions (2FIs) between cytokines for each 7-day interval. The *x* and *y* axis are the cytokine concentration (ng/ml) while the colorbar shows square root cell counts. (A) 2FIs for proT, CD4ISP, and early DP cells from day 0-14. (B) 2FIs for CD4ISP, early and late DP, and CD8ISP cells from day 14-35. No 2FIs were detected during day 35-42. RSM was constructed using n=3 pooled UCB donors.

### SUPPLEMENTAL TABLES

**Supplemental Table 1 (Related to Figure 1).** Factor concentrations for screening experiments.

| | Scaled factor level | | | |
|---|---|---|---|---|
| **Factor** | **-1** | **0** | **1** | |
| SCF | 1.5 | 15 | 150 | ng/ml |
| Flt3L | 1.5 | 15 | 150 | ng/ml |
| TPO | 1.5 | 15 | 150 | ng/ml |
| IL-1a | 0.01 | 0.1 | 1 | ng/ml |
| IL-1b | 0.01 | 0.1 | 1 | ng/ml |
| IL-3 | 0.25 | 2.5 | 25 | ng/ml |
| IL-6 | 0.25 | 2.5 | 25 | ng/ml |
| IL-7 | 1.5 | 15 | 150 | ng/ml |
| IL-15 | 0.25 | 2.5 | 25 | ng/ml |
| IL-18 | 0.25 | 2.5 | 25 | ng/ml |
| IL-21 | 0.25 | 2.5 | 25 | ng/ml |
| IFNγ | 0.25 | 2.5 | 25 | ng/ml |
| TNFa | 0.25 | 2.5 | 25 | ng/ml |
| RANTES | 0.25 | 2.5 | 25 | ng/ml |
| CXCL12 | 0.25 | 2.5 | 25 | ng/ml |

**Supplemental Table 2 (Related to Figure 1).** Linear regression model terms for total cells from day 0-7. Provided as raw values at the default precision in JMP 13 software that was used for regression.

| Term | Scaled Estimate | Std Error | t Ratio | Prob>\|t\| |
|---|---|---|---|---|
| Intercept | 78.5548504 | 4.15784548 | 18.8931625 | 1.17E-14 |
| SCF | 31.3582336 | 1.20371042 | 26.0513102 | 1.79E-17 |
| Flt3L | 2.34525045 | 1.20371042 | 1.94835104 | 0.06486147 |
| IL-1a | 2.54866038 | 1.20371042 | 2.11733681 | 0.04633552 |
| IL-1b | 1.112316 | 1.20371042 | 0.92407275 | 0.36594547 |
| IL-3 | 21.1497617 | 1.20371042 | 17.5704732 | 4.91E-14 |
| IL-7 | 2.96704437 | 1.20371042 | 2.46491541 | 0.02241018 |
| IL-18 | 2.16448163 | 1.20371042 | 1.7981747 | 0.08654215 |
| IL-21 | -7.1015991 | 1.20371042 | -5.8997571 | 7.42E-06 |
| IFNy | -1.8106322 | 1.20371042 | -1.5042091 | 0.14741957 |
| TNFa | 23.539892 | 1.20371042 | 19.5561088 | 5.87E-15 |
| RANTES | 0.16204596 | 1.20371042 | 0.13462205 | 0.89419264 |
| SCF*Flt3L | 7.10842437 | 1.6480454 | 4.31324549 | 0.00030721 |
| SCF*TNFa | -8.3335326 | 1.56078911 | -5.3393072 | 2.70E-05 |
| Flt3L*IL-1a | 10.7766857 | 1.69421559 | 6.36087034 | 2.63E-06 |
| IL-1b*IL-18 | -4.7305962 | 1.39240836 | -3.3974202 | 0.00271445 |
| IL-1b*TNFa | -9.4410142 | 1.36780148 | -6.9023278 | 8.06E-07 |
| IL-1b*RANTES | -4.4898134 | 1.60067235 | -2.8049547 | 0.01061109 |
| IL-3*IFNy | -2.5080706 | 1.71421754 | -1.4630994 | 0.15824749 |
| IL-3*RANTES | -3.6262683 | 1.70444908 | -2.127531 | 0.04538772 |
| IL-7*IL-7 | -17.328283 | 4.41783964 | -3.9223431 | 0.0007821 |

**Supplemental Table 3 (Related to Figure 1).** Linear regression model terms for CD7$^+$ cells from day 0-7. Provided as raw values at the default precision in JMP 13 software that was used for regression.

| Term | Scaled Estimate | Std Error | t Ratio | Prob>\|t\| |
|---|---|---|---|---|
| Intercept | 38.2354304 | 2.28993593 | 16.6971617 | 1.03E-14 |
| SCF | 3.46255372 | 0.70658425 | 4.90041168 | 5.35E-05 |
| Flt3L | 0.7221513 | 0.70658425 | 1.0220314 | 0.31695825 |
| IL-1a | 0.93794616 | 0.70658425 | 1.32743712 | 0.19685369 |
| IL-3 | 6.96508011 | 0.70658425 | 9.85739505 | 6.50E-10 |
| IL-7 | 2.92510744 | 0.70658425 | 4.13978578 | 0.00036989 |
| IL-15 | -1.4546059 | 0.70658425 | -2.0586447 | 0.05054079 |
| IL-18 | 0.59658713 | 0.70658425 | 0.84432554 | 0.40682544 |
| IFNγ | -1.4251344 | 0.70658425 | -2.0169349 | 0.05502206 |
| TNFa | 17.1896859 | 0.70658425 | 24.3278644 | 2.01E-18 |
| RANTES | -0.7447359 | 0.70658425 | -1.0539945 | 0.3023814 |
| SCF*IL-7 | -6.1424113 | 0.95590818 | -6.4257336 | 1.21E-06 |
| SCF*IFNγ | -5.1511001 | 0.79602838 | -6.4710006 | 1.08E-06 |
| SCF*TNFa | -5.8116604 | 0.78933744 | -7.3627071 | 1.33E-07 |
| Flt3L*IFNγ | -2.8996081 | 0.85006097 | -3.411059 | 0.00229435 |
| IL-1a*IL-18 | -2.0964354 | 0.81809125 | -2.5625936 | 0.01708609 |
| IL-1a*RANTES | 4.4779813 | 0.85660531 | 5.22758994 | 2.34E-05 |
| IFNγ*IFNγ | -9.5840397 | 2.41951521 | -3.9611405 | 0.00058124 |

**Supplemental Table 4 (Related to Figure 1).** Linear regression model terms for $CD7^{+}CD5^{+}$ cells from day 0-7. Provided as raw values at the default precision in JMP 13 software that was used for regression.

| Term | Scaled Estimate | Std Error | t Ratio | Prob>\|t\| |
|---|---|---|---|---|
| Intercept | 88.2119145 | 7.44984267 | 11.8407755 | 1.65E-11 |
| Block[1] | 12.2212261 | 2.32187531 | 5.26351523 | 2.14E-05 |
| Block[2] | -12.221226 | 2.32187531 | -5.2635152 | 2.14E-05 |
| SCF | 11.8872455 | 2.09706254 | 5.66852217 | 7.74E-06 |
| TPO | -3.5716481 | 2.09706254 | -1.7031672 | 0.10145437 |
| IL-3 | 16.6548214 | 2.09706254 | 7.94197648 | 3.59E-08 |
| IL-15 | -4.9546677 | 2.09706254 | -2.3626705 | 0.02659002 |
| IL-21 | 3.75240108 | 2.09706254 | 1.7893606 | 0.08618547 |
| IFNγ | -10.561468 | 2.09706254 | -5.036315 | 3.79E-05 |
| TNFa | 17.4650403 | 2.09706254 | 8.32833548 | 1.54E-08 |
| CXCL12 | 0.98962735 | 2.09706254 | 0.47191123 | 0.64125256 |
| SCF*IFNγ | -9.6348035 | 2.42995194 | -3.9650181 | 0.00057558 |
| TPO*IFNγ | -9.791543 | 2.52912493 | -3.8715142 | 0.00072856 |
| IL-3*IL-15 | -9.9531221 | 2.5795491 | -3.8584736 | 0.00075287 |
| IL-3*IL-21 | 10.5685464 | 2.41458907 | 4.37695445 | 0.00020254 |
| IL-3*CXCL12 | -10.476515 | 2.66295626 | -3.934167 | 0.00062217 |
| IL-15*IL-21 | -5.1613043 | 2.53626398 | -2.0350028 | 0.05303928 |
| IFNγ*TNFa | 7.59653929 | 2.67356555 | 2.84135143 | 0.0090177 |
| IL-15*IL-15 | -52.338332 | 7.9333962 | -6.5972165 | 8.00E-07 |

**Supplemental Table 5 (Related to Figure 1).** Linear regression model terms for total cells from day 7-14. Provided as raw values at the default precision in JMP 13 software that was used for regression.

| Term | Scaled Estimate | Std Error | t Ratio | Prob>\|t\| |
|---|---|---|---|---|
| Intercept | -0.4294464 | 1.05235746 | -0.4080804 | 0.68655584 |
| Block[1] | -5.0276265 | 0.31540451 | -15.940249 | 6.16E-15 |
| Block[2] | 5.02762648 | 0.31540451 | 15.9402491 | 6.16E-15 |
| SCF | 7.19690786 | 0.32688977 | 22.0163141 | 2.45E-18 |
| Flt3L | 1.19713814 | 0.32688977 | 3.6622074 | 0.00112089 |
| IL-1a | 1.37285996 | 0.32688977 | 4.1997642 | 0.00027722 |
| IL-3 | 3.17619827 | 0.32688977 | 9.71641988 | 3.85E-10 |
| IL-6 | -1.4756853 | 0.32688977 | -4.5143208 | 0.00012114 |
| IL-7 | 1.65815678 | 0.32688977 | 5.07252576 | 2.78E-05 |
| IL-18 | 0.55791221 | 0.32688977 | 1.70672886 | 0.09979047 |
| IL-21 | -0.759394 | 0.32688977 | -2.3230888 | 0.02826392 |
| IFNγ | -0.7815329 | 0.32688977 | -2.3908148 | 0.02434207 |
| TNFa | 2.91092812 | 0.32688977 | 8.90492264 | 2.24E-09 |
| RANTES | -1.454269 | 0.32688977 | -4.4488054 | 0.00014398 |
| Flt3L*RANTES | 0.76822075 | 0.342659 | 2.24193955 | 0.03371702 |
| IL-3*RANTES | 0.58978318 | 0.342659 | 1.72119568 | 0.09709567 |
| IL-21*IL-21 | 4.54250539 | 1.11120245 | 4.08791882 | 0.00037159 |

**Supplemental Table 6 (Related to Figure 1).** Linear regression model terms for $CD7^{+}$ cells from day 7-14. Provided as raw values at the default precision in JMP 13 software that was used for regression.

| Term | Scaled Estimate | Std Error | t Ratio | Prob>\|t\| |
|---|---|---|---|---|
| Intercept | 8.46044005 | 1.27962456 | 6.61165805 | 1.94E-06 |
| Block[1] | -5.0928099 | 0.43910176 | -11.598245 | 2.48E-10 |
| Block[2] | 5.09280988 | 0.43910176 | 11.5982451 | 2.48E-10 |
| SCF | 3.79869897 | 0.39057606 | 9.72588788 | 5.04E-09 |
| Flt3L | 1.13930796 | 0.39057606 | 2.91699383 | 0.0085242 |
| TPO | 0.79111712 | 0.39057606 | 2.0255136 | 0.05636689 |
| IL-1a | 1.67013143 | 0.39057606 | 4.27607218 | 0.00036892 |
| IL-1b | 1.94766572 | 0.39057606 | 4.986649 | 7.09E-05 |
| IL-3 | 2.81688755 | 0.39057606 | 7.21213572 | 5.56E-07 |
| IL-6 | -0.8633749 | 0.39057606 | -2.2105167 | 0.0388855 |
| IL-7 | 3.73510864 | 0.39057606 | 9.56307625 | 6.67E-09 |
| IL-15 | 0.55677285 | 0.39057606 | 1.4255171 | 0.16942901 |
| IL-21 | -0.4414777 | 0.39057606 | -1.1303246 | 0.2717132 |
| IFNγ | -1.5284321 | 0.39057606 | -3.9132765 | 0.00086162 |
| TNFa | 4.87670456 | 0.39057606 | 12.485928 | 6.72E-11 |
| RANTES | -0.9652458 | 0.39057606 | -2.4713389 | 0.02257089 |
| SCF*TNFa | -0.9089791 | 0.47216179 | -1.9251432 | 0.06855202 |
| TPO*IL-1b | 3.23140275 | 0.50087236 | 6.4515493 | 2.72E-06 |
| TPO*IL-21 | -1.8752665 | 0.48234336 | -3.8878249 | 0.00091441 |
| IL-1a*TNFa | 2.00931716 | 0.44444679 | 4.52093976 | 0.00020836 |
| IL-3*IFNγ | 0.86804295 | 0.55702447 | 1.55835695 | 0.13483258 |
| IL-7*TNFa | 2.1101213 | 0.51227722 | 4.11910037 | 0.0005325 |
| IL-3*IL-3 | -3.7070962 | 1.3534027 | -2.7390932 | 0.01264601 |

**Supplemental Table 7 (Related to Figure 1).** Linear regression model terms for $CD7^{+}CD5^{+}$ cells from day 7-14. Provided as raw values at the default precision in JMP 13 software that was used for regression.

| Term | Scaled Estimate | Std Error | t Ratio | Prob>\|t\| |
|---|---|---|---|---|
| Intercept | 1.41785632 | 0.17358573 | 8.16804634 | 2.75E-07 |
| Block[1] | -0.5518726 | 0.05768997 | -9.5661803 | 2.95E-08 |
| Block[2] | 0.55187261 | 0.05768997 | 9.5661803 | 2.95E-08 |
| SCF | 0.69330822 | 0.04658126 | 14.8838456 | 3.50E-11 |
| Flt3L | 0.15000886 | 0.04658126 | 3.22036967 | 0.00502245 |
| TPO | 0.09709329 | 0.04658126 | 2.08438542 | 0.05252144 |
| IL-1a | 0.15465599 | 0.04658126 | 3.32013345 | 0.00405013 |
| IL-1b | 0.08215625 | 0.04658126 | 1.76371908 | 0.0957415 |
| IL-3 | 0.33033842 | 0.04658126 | 7.0916597 | 1.81E-06 |
| IL-6 | -0.103375 | 0.04658126 | -2.2192409 | 0.04036517 |
| IL-7 | 0.45819466 | 0.04658126 | 9.83645997 | 1.97E-08 |
| IL-15 | -0.1457697 | 0.04658126 | -3.129364 | 0.00610763 |
| IL-18 | -0.1329391 | 0.04658126 | -2.8539173 | 0.01098298 |
| IL-21 | -0.1124967 | 0.04658126 | -2.4150635 | 0.02728165 |
| IFNγ | -0.5120009 | 0.04658126 | -10.991565 | 3.81E-09 |
| TNFa | 0.58937963 | 0.04658126 | 12.6527209 | 4.46E-10 |
| RANTES | -0.1170682 | 0.04658126 | -2.513204 | 0.02233552 |
| CXCL12 | 0.00768674 | 0.04658126 | 0.16501783 | 0.87087631 |
| IL-1a*IL-7 | 0.33797203 | 0.07599821 | 4.44710524 | 0.0003536 |
| IL-1b*TNFa | -0.2325363 | 0.05320178 | -4.3708372 | 0.00041646 |
| IL-6*IFNγ | 0.162921 | 0.06287869 | 2.59103686 | 0.01902942 |
| IL-7*TNFa | 0.25344169 | 0.05413476 | 4.68168096 | 0.00021438 |
| IL-18*TNFa | 0.17736364 | 0.05940917 | 2.98545924 | 0.00830821 |
| TNFa*CXCL12 | -0.3137906 | 0.05479505 | -5.7266228 | 2.47E-05 |
| IL-1b*IL-1b | -0.4097009 | 0.21091751 | -1.9424697 | 0.06883406 |
| IL-18*IL-18 | -1.6892753 | 0.20882721 | -8.0893451 | 3.14E-07 |

**Supplemental Table 8 (Related to Supplemental Figure 1).** Factor concentrations for screening validation experiments.

| | Concentration (ng/ml) | | |
|---|---|---|---|
| **Factor** | **-1** | **0** | **1** |
| SCF | 3 | 30 | 300 |
| Flt3L | 3 | 30 | 300 |
| TPO | 3 | 30 | 300 |
| IL-1a | 0.5 | 5 | 50 |
| IL-1b | 0.5 | 5 | 50 |
| IL-3 | 1 | 10 | 100 |
| IL-7 | 3 | 30 | 300 |
| IL-21 | 1 | 10 | 100 |
| TNFa | 1 | 10 | 100 |
| CXCL12 | 3 | 30 | 300 |

**Supplemental Table 9 (Related to Supplemental Figure 1).** Linear regression model terms for total cells from day 0-7. Provided as raw values at the default precision in JMP 13 software that was used for regression.

| Term | Scaled Estimate | Std Error | t Ratio | Prob>\|t\| |
|---|---|---|---|---|
| Intercept | 421.7535 | 15.79248 | 26.70597 | 1.57E-16 |
| SCF | 49.50128 | 6.087245 | 8.131967 | 1.31E-07 |
| Flt3L | 19.94848 | 6.087245 | 3.277095 | 0.003965 |
| IL-1a | -23.8007 | 6.087245 | -3.90992 | 0.000941 |
| IL-1b | 23.58617 | 6.087245 | 3.874688 | 0.00102 |
| IL-3 | 3.677086 | 6.087245 | 0.604064 | 0.552946 |
| IL-7 | -12.4957 | 6.087245 | -2.05277 | 0.054127 |
| IL-21 | -26.9593 | 6.087245 | -4.42882 | 0.000288 |
| TNFa | 31.11326 | 6.087245 | 5.111221 | 6.21E-05 |
| CXCL12 | 9.255202 | 6.087245 | 1.520425 | 0.144872 |
| Flt3L*Flt3L | -96.1159 | 21.97905 | -4.37307 | 0.000327 |
| IL-1a*IL-1a | 63.68215 | 22.61573 | 2.815835 | 0.011036 |
| TNFa*TNFa | -96.2695 | 21.99432 | -4.37702 | 0.000324 |
| Flt3L*IL-3 | 29.90263 | 7.14127 | 4.187299 | 0.0005 |
| Flt3L*CXCL12 | 15.38151 | 6.572035 | 2.340448 | 0.030323 |
| IL-1a*IL-3 | -29.4113 | 6.606348 | -4.45197 | 0.000273 |
| IL-21*TNFa | 13.91181 | 7.072824 | 1.966939 | 0.063968 |

**Supplemental Table 10 (Related to Supplemental Figure 1).** Linear regression model terms for $CD7^+$ cells from day 0-7. Provided as raw values at the default precision in JMP 13 software that was used for regression.

| Term | Scaled Estimate | Std Error | t Ratio | Prob>\|t\| |
|---|---|---|---|---|
| Intercept | 324.0167 | 11.23244 | 28.84652 | 1.6E-16 |
| SCF | 23.20677 | 4.231553 | 5.48422 | 3.3E-05 |
| Flt3L | 18.73727 | 4.231553 | 4.427989 | 0.000325 |
| IL-1a | -17.9357 | 4.231553 | -4.23856 | 0.000494 |
| IL-1b | 19.39555 | 4.231553 | 4.583554 | 0.00023 |
| IL-3 | 4.937447 | 4.231553 | 1.166817 | 0.258511 |
| IL-7 | -5.87237 | 4.231553 | -1.38776 | 0.182152 |
| IL-21 | -7.96603 | 4.231553 | -1.88253 | 0.076031 |
| TNFa | 34.63903 | 4.231553 | 8.18589 | 1.76E-07 |
| CXCL12 | 8.542467 | 4.231553 | 2.018754 | 0.058663 |
| Flt3L*Flt3L | -37.2207 | 15.33467 | -2.42723 | 0.025934 |
| IL-1a*IL-1a | 50.16291 | 16.83598 | 2.979506 | 0.008035 |
| IL-3*IL-3 | -25.7467 | 14.75829 | -1.74456 | 0.098113 |
| TNFa*TNFa | -92.0613 | 17.63344 | -5.22083 | 5.77E-05 |
| SCF*IL-3 | 7.680827 | 5.343498 | 1.437416 | 0.167757 |
| Flt3L*IL-3 | 16.3621 | 4.998596 | 3.273339 | 0.004223 |
| Flt3L*CXCL12 | 13.04087 | 4.75419 | 2.743027 | 0.01337 |
| IL-1a*IL-3 | -26.431 | 4.705941 | -5.61652 | 2.49E-05 |

**Supplemental Table 11 (Related to Supplemental Figure 1).** Linear regression model terms for $CD7^{+}CD5^{+}$ cells from day 0-7. Provided as raw values at the default precision in JMP 13 software that was used for regression.

| Term | Scaled Estimate | Std Error | t Ratio | Prob>\|t\| |
|---|---|---|---|---|
| Intercept | 120.3282 | 4.597319 | 26.17357 | 1.25E-12 |
| Block[1] | -13.3621 | 4.049695 | -3.29953 | 0.005753 |
| Block[2] | 13.36209 | 4.049695 | 3.29953 | 0.005753 |
| SCF | 16.45338 | 1.729571 | 9.512984 | 3.2E-07 |
| Flt3L | 7.705694 | 1.729571 | 4.455263 | 0.000648 |
| IL-1a | -10.2907 | 1.729571 | -5.94986 | 4.83E-05 |
| IL-1b | 9.151321 | 1.729571 | 5.291093 | 0.000146 |
| IL-3 | 4.077907 | 1.729571 | 2.357756 | 0.03472 |
| IL-7 | -3.96261 | 1.729571 | -2.29109 | 0.039303 |
| IL-21 | -4.48671 | 1.729571 | -2.59412 | 0.022253 |
| TNFa | 1.505027 | 1.729571 | 0.870173 | 0.399988 |
| CXCL12 | 3.460036 | 1.729571 | 2.000517 | 0.066779 |
| Flt3L*Flt3L | -13.4037 | 15.08061 | -0.8888 | 0.390259 |
| IL-7*IL-7 | -33.4941 | 14.42399 | -2.32211 | 0.037103 |
| SCF*IL-1b | -6.12621 | 2.798603 | -2.18902 | 0.047444 |
| SCF*IL-21 | -10.3563 | 5.692225 | -1.81937 | 0.091953 |
| SCF*TNFa | -9.03616 | 2.24282 | -4.02893 | 0.001432 |
| IL-1a*IL-1b | 12.40104 | 2.364721 | 5.24419 | 0.000158 |
| IL-1a*IL-3 | -7.47573 | 2.702475 | -2.76625 | 0.016032 |
| IL-1a*TNFa | -14.3917 | 7.582211 | -1.89809 | 0.080111 |
| IL-1b*IL-3 | -1.94854 | 3.671769 | -0.53068 | 0.604588 |
| IL-1b*IL-7 | 14.24745 | 7.460918 | 1.909612 | 0.078498 |
| IL-1b*CXCL12 | 2.759172 | 3.646236 | 0.756718 | 0.462713 |
| IL-3*TNFa | -1.55565 | 2.754867 | -0.56469 | 0.581897 |

**Supplemental Table 12 (Related to Supplemental Figure 1).** Linear regression model terms for total cells from day 7-14. Provided as raw values at the default precision in JMP 13 software that was used for regression.

| Term | Scaled Estimate | Std Error | t Ratio | Prob>\|t\| |
|---|---|---|---|---|
| Intercept | 237.825 | 3.770277 | 63.07892 | 1.42E-22 |
| Block[1] | 12.96798 | 1.934785 | 6.702544 | 2.76E-06 |
| Block[2] | -12.968 | 1.934785 | -6.70254 | 2.76E-06 |
| SCF | 20.31032 | 1.44464 | 14.05909 | 3.79E-11 |
| IL-1b | 0.319537 | 1.44464 | 0.221188 | 0.827436 |
| IL-3 | 8.790551 | 1.44464 | 6.084943 | 9.47E-06 |
| IL-7 | 72.43007 | 1.44464 | 50.13711 | 8.64E-21 |
| IL-21 | -10.2499 | 1.44464 | -7.09516 | 1.3E-06 |
| TNFa | 15.64402 | 1.44464 | 10.82901 | 2.59E-09 |
| CXCL12 | 4.857228 | 1.44464 | 3.362241 | 0.00347 |
| IL-1b*IL-3 | -2.55604 | 1.807759 | -1.41393 | 0.174447 |
| IL-1b*CXCL12 | -9.80432 | 1.805492 | -5.43028 | 3.69E-05 |
| IL-3*CXCL12 | -3.30956 | 1.874916 | -1.76518 | 0.094493 |
| IL-7*IL-21 | -7.84978 | 1.619706 | -4.84643 | 0.00013 |
| IL-7*TNFa | 15.34255 | 1.820535 | 8.427498 | 1.16E-07 |
| IL-7*CXCL12 | 2.905552 | 1.626214 | 1.786698 | 0.09084 |
| TNFa*CXCL12 | 3.862878 | 1.70058 | 2.271506 | 0.03562 |
| SCF*SCF | -19.3747 | 5.094722 | -3.8029 | 0.001303 |
| IL-7*IL-7 | -65.1195 | 4.913722 | -13.2526 | 1E-10 |

**Supplemental Table 13 (Related to Supplemental Figure 1).** Linear regression model terms for CD7$^+$ cells from day 7-14. Provided as raw values at the default precision in JMP 13 software that was used for regression.

| Term | Scaled Estimate | Std Error | t Ratio | Prob>\|t\| |
|---|---|---|---|---|
| Intercept | 213.0429 | 4.340803 | 49.07915 | 1.77E-21 |
| Block[1] | 8.018208 | 2.394469 | 3.348638 | 0.003374 |
| Block[2] | -8.01821 | 2.394469 | -3.34864 | 0.003374 |
| SCF | 8.842854 | 1.660564 | 5.325211 | 3.87E-05 |
| IL-3 | 7.327362 | 1.660564 | 4.412574 | 0.000299 |
| IL-7 | 83.47695 | 1.660564 | 50.27024 | 1.12E-21 |
| IL-21 | -10.2832 | 1.660564 | -6.1926 | 5.98E-06 |
| TNFa | 12.86034 | 1.660564 | 7.744562 | 2.7E-07 |
| CXCL12 | 4.224562 | 1.660564 | 2.544052 | 0.019807 |
| SCF*IL-7 | 7.063645 | 2.466983 | 2.863273 | 0.009949 |
| SCF*TNFa | -3.81721 | 1.876099 | -2.03465 | 0.056084 |
| SCF*CXCL12 | -5.59685 | 2.105053 | -2.65877 | 0.015506 |
| IL-7*IL-21 | -13.3368 | 1.967794 | -6.77753 | 1.79E-06 |
| IL-7*TNFa | 13.38199 | 2.162249 | 6.188922 | 6.03E-06 |
| IL-7*CXCL12 | 6.130731 | 1.926027 | 3.183098 | 0.004896 |
| IL-3*IL-3 | -15.4358 | 9.207936 | -1.67636 | 0.110045 |
| IL-7*IL-7 | -55.4034 | 6.933754 | -7.99039 | 1.71E-07 |
| TNFa*TNFa | -23.8174 | 7.084963 | -3.36168 | 0.003276 |

**Supplemental Table 14 (Related to Supplemental Figure 1).** Linear regression model terms for $CD7^{+}CD5^{+}$ cells from day 7-14. Provided as raw values at the default precision in JMP 13 software that was used for regression.

| Term | Scaled Estimate | Std Error | t Ratio | Prob>\|t\| |
|---|---|---|---|---|
| Intercept | 237.825 | 3.770277 | 63.07892 | 1.42E-22 |
| Block[1] | 12.96798 | 1.934785 | 6.702544 | 2.76E-06 |
| Block[2] | -12.968 | 1.934785 | -6.70254 | 2.76E-06 |
| SCF | 20.31032 | 1.44464 | 14.05909 | 3.79E-11 |
| IL-1b | 0.319537 | 1.44464 | 0.221188 | 0.827436 |
| IL-3 | 8.790551 | 1.44464 | 6.084943 | 9.47E-06 |
| IL-7 | 72.43007 | 1.44464 | 50.13711 | 8.64E-21 |
| IL-21 | -10.2499 | 1.44464 | -7.09516 | 1.3E-06 |
| TNFa | 15.64402 | 1.44464 | 10.82901 | 2.59E-09 |
| CXCL12 | 4.857228 | 1.44464 | 3.362241 | 0.00347 |
| IL-1b*IL-3 | -2.55604 | 1.807759 | -1.41393 | 0.174447 |
| IL-1b*CXCL12 | -9.80432 | 1.805492 | -5.43028 | 3.69E-05 |
| IL-3*CXCL12 | -3.30956 | 1.874916 | -1.76518 | 0.094493 |
| IL-7*IL-21 | -7.84978 | 1.619706 | -4.84643 | 0.00013 |
| IL-7*TNFa | 15.34255 | 1.820535 | 8.427498 | 1.16E-07 |
| IL-7*CXCL12 | 2.905552 | 1.626214 | 1.786698 | 0.09084 |
| TNFa*CXCL12 | 3.862878 | 1.70058 | 2.271506 | 0.03562 |
| SCF*SCF | -19.3747 | 5.094722 | -3.8029 | 0.001303 |
| IL-7*IL-7 | -65.1195 | 4.913722 | -13.2526 | 1E-10 |

**Supplemental Table 15 (Related to Figure 4).** Factor concentrations for RSM experiments.

| | Scaled factor level | | | | | |
|---|---|---|---|---|---|---|
| **Factor** | **-2.366** | **-1** | **0** | **1** | **2.366** | |
| SCF | 0.52 | 2.86 | 10.00 | 35.00 | 193.76 | ng/ml |
| Flt3L | 0.52 | 2.86 | 10.00 | 35.00 | 193.76 | ng/ml |
| IL-3 | 0.05 | 0.29 | 1.00 | 3.50 | 19.38 | ng/ml |
| IL-7 | 0.52 | 2.86 | 10.00 | 35.00 | 193.76 | ng/ml |
| TNFa | 0.03 | 0.14 | 0.50 | 1.75 | 9.69 | ng/ml |
| CXCL12 | 0.52 | 2.86 | 10.00 | 35.00 | 193.76 | ng/ml |

**Supplemental Figure 16 (Related to Figure 4).** Linear regression model terms for ProT-cells from day 0-7. Provided as raw values at the default precision in JMP 14 software that was used for regression.

| Term | Estimate | Std Error | t Ratio | Prob>\|t\| |
|---|---|---|---|---|
| Intercept | 71.6797471 | 1.67416813 | 42.82 | <.0001 |
| SCF | 9.9138704 | 1.85717241 | 5.34 | <.0001 |
| IL-3 | 15.8415123 | 1.85717241 | 8.53 | <.0001 |
| TNFa | 32.033221 | 1.85717241 | 17.25 | <.0001 |
| SCF*TNFa | 7.19163125 | 2.15783934 | 3.33 | 0.0017 |
| IL-3*TNFa | 10.4289214 | 2.15783934 | 4.83 | <.0001 |
| Block[1] | 7.6578892 | 2.29917013 | 3.33 | 0.0017 |
| Block[2] | -0.2290307 | 2.32742038 | -0.1 | 0.922 |

**Supplemental Figure 17 (Related to Figure 4).** Linear regression model terms for CD4ISP cells from day 0-7. Provided as raw values at the default precision in JMP 14 software that was used for regression.

| Term | Estimate | Std Error | t Ratio | Prob>\|t\| |
|---|---|---|---|---|
| Intercept | 103.61767 | 2.0413071 | 50.76 | <.0001 |
| SCF | 16.8207395 | 2.26444355 | 7.43 | <.0001 |
| IL-3 | 13.8267354 | 2.26444355 | 6.11 | <.0001 |
| TNFa | 17.5943395 | 2.26444355 | 7.77 | <.0001 |
| SCF*TNFa | 4.70545012 | 2.63104565 | 1.79 | 0.0803 |
| IL-3*TNFa | 2.48957783 | 2.63104565 | 0.95 | 0.349 |
| Block[1] | 14.2788352 | 2.80336975 | 5.09 | <.0001 |
| Block[2] | 1.7624824 | 2.83781518 | 0.62 | 0.5376 |

**Supplemental Figure 18 (Related to Figure 4).** Linear regression model terms for early DP cells from day 0-7. Provided as raw values at the default precision in JMP 14 software that was used for regression.

| Term | Estimate | Std Error | t Ratio | Prob>\|t\| |
|---|---|---|---|---|
| Intercept | 23.5976571 | 0.51419461 | 45.89 | <.0001 |
| SCF | 1.23598546 | 0.57040152 | 2.17 | 0.0355 |
| IL-3 | 2.32434246 | 0.57040152 | 4.07 | 0.0002 |
| TNFa | 3.15202229 | 0.57040152 | 5.53 | <.0001 |
| SCF*TNFa | 0.1023632 | 0.66274667 | 0.15 | 0.8779 |
| IL-3*TNFa | 0.38530373 | 0.66274667 | 0.58 | 0.5638 |
| Block[1] | 4.96773292 | 0.70615421 | 7.03 | <.0001 |
| Block[2] | 0.5066865 | 0.71483084 | 0.71 | 0.482 |

**Supplemental Figure 19 (Related to Figure 4).** Linear regression model terms for proT-cells from day 7-14. Provided as raw values at the default precision in JMP 14 software that was used for regression.

| Term | Estimate | Std Error | t Ratio | Prob>\|t\| |
|---|---|---|---|---|
| Intercept | 291.628979 | 6.14680328 | 47.44 | <.0001 |
| SCF | 17.6115579 | 3.6017748 | 4.89 | <.0001 |
| Flt3L | -4.0484279 | 3.6017748 | -1.12 | 0.2677 |
| IL-7 | 174.729196 | 5.32166436 | 32.83 | <.0001 |
| TNFa | -0.0298286 | 3.6017748 | -0.01 | 0.9934 |
| CXCL12 | 12.0255116 | 3.6017748 | 3.34 | 0.0018 |
| TNFa*CXCL12 | -26.458075 | 4.18488415 | -6.32 | <.0001 |
| SCF*SCF | -4.6428883 | 3.06379363 | -1.52 | 0.1375 |
| Flt3L*Flt3L | -13.788429 | 3.06379363 | -4.5 | <.0001 |
| IL-7*IL-7 | -12.485266 | 3.06379363 | -4.08 | 0.0002 |
| CXCL12*CXCL12 | -8.8235862 | 3.06379363 | -2.88 | 0.0064 |
| Block[1] | 39.2710377 | 4.45904662 | 8.81 | <.0001 |
| Block[2] | 5.06404619 | 4.52376056 | 1.12 | 0.2696 |
| IL-7*IL-7*IL-7 | -21.397621 | 1.78672811 | -11.98 | <.0001 |

**Supplemental Figure 20 (Related to Figure 4).** Linear regression model terms for CD4ISP cells from day 7-14. Provided as raw values at the default precision in JMP 14 software that was used for regression.

| Term | Estimate | Std Error | t Ratio | Prob>\|t\| |
|---|---|---|---|---|
| Intercept | 133.739148 | 4.0060544 | 33.38 | <.0001 |
| SCF | 24.729668 | 2.3473837 | 10.53 | <.0001 |
| Flt3L | -2.3644042 | 2.3473837 | -1.01 | 0.3199 |
| IL-7 | 23.7706908 | 3.46828684 | 6.85 | <.0001 |
| TNFa | 5.02661963 | 2.3473837 | 2.14 | 0.0384 |
| CXCL12 | 5.12097499 | 2.3473837 | 2.18 | 0.0351 |
| TNFa*CXCL12 | -5.9615814 | 2.72741339 | -2.19 | 0.0347 |
| SCF*SCF | -3.0420872 | 1.99676538 | -1.52 | 0.1355 |
| Flt3L*Flt3L | -7.4211583 | 1.99676538 | -3.72 | 0.0006 |
| IL-7*IL-7 | -9.420782 | 1.99676538 | -4.72 | <.0001 |
| CXCL12*CXCL12 | -5.5856195 | 1.99676538 | -2.8 | 0.0079 |
| Block[1] | 19.3786765 | 2.90609323 | 6.67 | <.0001 |
| Block[2] | 0.11294902 | 2.94826922 | 0.04 | 0.9696 |
| IL-7*IL-7*IL-7 | -2.4283895 | 1.16446382 | -2.09 | 0.0435 |

**Supplemental Figure 21 (Related to Figure 4).** Linear regression model terms for early DP cells from day 7-14. Provided as raw values at the default precision in JMP 14 software that was used for regression.

| Term | Estimate | Std Error | t Ratio | Prob>\|t\| |
|---|---|---|---|---|
| Intercept | 27.4192258 | 1.02921097 | 26.64 | <.0001 |
| SCF | 4.99336479 | 0.60307545 | 8.28 | <.0001 |
| Flt3L | -1.049385 | 0.60307545 | -1.74 | 0.0895 |
| IL-7 | 9.67417562 | 0.89105102 | 10.86 | <.0001 |
| TNFa | 0.45534857 | 0.60307545 | 0.76 | 0.4546 |
| CXCL12 | 1.29563789 | 0.60307545 | 2.15 | 0.0378 |
| TNFa*CXCL12 | -2.2437249 | 0.70071035 | -3.2 | 0.0027 |
| SCF*SCF | -0.1581914 | 0.51299673 | -0.31 | 0.7594 |
| Flt3L*Flt3L | -1.3805223 | 0.51299673 | -2.69 | 0.0103 |
| IL-7*IL-7 | -1.5753906 | 0.51299673 | -3.07 | 0.0038 |
| CXCL12*CXCL12 | -1.3714581 | 0.51299673 | -2.67 | 0.0108 |
| Block[1] | 3.71456732 | 0.74661568 | 4.98 | <.0001 |
| Block[2] | 0.35401123 | 0.75745127 | 0.47 | 0.6428 |
| IL-7*IL-7*IL-7 | -1.1331093 | 0.29916691 | -3.79 | 0.0005 |

**Supplemental Figure 22 (Related to Figure 4).** Linear regression model terms for CD4ISP cells from day 14-21. Provided as raw values at the default precision in JMP 14 software that was used for regression.

| Term | Estimate | Std Error | t Ratio | Prob>\|t\| |
|---|---|---|---|---|
| Intercept | 120.237143 | 2.96960579 | 40.49 | <.0001 |
| SCF | 27.0731414 | 2.63666224 | 10.27 | <.0001 |
| Flt3L | -1.44686 | 2.63666224 | -0.55 | 0.5862 |
| IL-3 | 3.88158144 | 2.63666224 | 1.47 | 0.1488 |
| IL-7 | 30.4154464 | 3.89569926 | 7.81 | <.0001 |
| TNFa | 4.03558101 | 2.63666224 | 1.53 | 0.1337 |
| CXCL12 | 3.03577086 | 2.63666224 | 1.15 | 0.2564 |
| Flt3L*IL-3 | -0.4743265 | 3.06352468 | -0.15 | 0.8777 |
| SCF*IL-7 | 2.81594141 | 3.06352468 | 0.92 | 0.3635 |
| TNFa*CXCL12 | -7.4490183 | 3.06352468 | -2.43 | 0.0196 |
| IL-7*IL-7 | -6.8304829 | 2.23572417 | -3.06 | 0.004 |
| Block[1] | 21.452036 | 3.26418535 | 6.57 | <.0001 |
| Block[2] | 3.90525978 | 3.30587154 | 1.18 | 0.2445 |
| IL-7*IL-7*IL-7 | -3.0163804 | 1.30796588 | -2.31 | 0.0264 |

**Supplemental Figure 23 (Related to Figure 4).** Linear regression model terms for early DP cells from day 14-21. Provided as raw values at the default precision in JMP 14 software that was used for regression.

| Term | Estimate | Std Error | t Ratio | Prob>\|t\| |
|---|---|---|---|---|
| Intercept | 52.9982187 | 1.2782537 | 41.46 | <.0001 |
| SCF | 10.4819647 | 1.13493962 | 9.24 | <.0001 |
| Flt3L | -1.0781755 | 1.13493962 | -0.95 | 0.3478 |
| IL-3 | 1.0539978 | 1.13493962 | 0.93 | 0.3586 |
| IL-7 | 16.5635165 | 1.67688655 | 9.88 | <.0001 |
| TNFa | 2.66784135 | 1.13493962 | 2.35 | 0.0238 |
| CXCL12 | 2.38344237 | 1.13493962 | 2.1 | 0.0421 |
| Flt3L*IL-3 | 0.43283012 | 1.31868068 | 0.33 | 0.7445 |
| SCF*IL-7 | 1.56543433 | 1.31868068 | 1.19 | 0.2422 |
| TNFa*CXCL12 | -2.4524409 | 1.31868068 | -1.86 | 0.0703 |
| IL-7*IL-7 | -2.3078868 | 0.9623576 | -2.4 | 0.0212 |
| Block[1] | 10.7118481 | 1.40505418 | 7.62 | <.0001 |
| Block[2] | 1.85193911 | 1.42299781 | 1.3 | 0.2006 |
| IL-7*IL-7*IL-7 | -1.8447401 | 0.56300814 | -3.28 | 0.0022 |

**Supplemental Figure 24 (Related to Figure 4).** Linear regression model terms for late DP cells from day 14-21. Provided as raw values at the default precision in JMP 14 software that was used for regression.

| Term | Estimate | Std Error | t Ratio | Prob>\|t\| |
|---|---|---|---|---|
| Intercept | 9.70567708 | 0.36147161 | 26.85 | <.0001 |
| SCF | 0.956464 | 0.32094446 | 2.98 | 0.0049 |
| Flt3L | -0.273237 | 0.32094446 | -0.85 | 0.3996 |
| IL-3 | 0.16991784 | 0.32094446 | 0.53 | 0.5994 |
| IL-7 | 3.62133847 | 0.4741992 | 7.64 | <.0001 |
| TNFa | -0.28819 | 0.32094446 | -0.9 | 0.3746 |
| CXCL12 | 0.2827937 | 0.32094446 | 0.88 | 0.3835 |
| Flt3L*IL-3 | -1.1503788 | 0.37290377 | -3.08 | 0.0037 |
| SCF*IL-7 | 1.03096742 | 0.37290377 | 2.76 | 0.0086 |
| TNFa*CXCL12 | -0.9698463 | 0.37290377 | -2.6 | 0.013 |
| IL-7*IL-7 | -0.7988181 | 0.27214077 | -2.94 | 0.0055 |
| Block[1] | 1.52363388 | 0.39732894 | 3.83 | 0.0004 |
| Block[2] | 0.42112253 | 0.40240314 | 1.05 | 0.3016 |
| IL-7*IL-7*IL-7 | -0.6375127 | 0.15921054 | -4 | 0.0003 |

**Supplemental Figure 25 (Related to Figure 4).** Linear regression model terms for CD8SP cells from day 14-21. Provided as raw values at the default precision in JMP 14 software that was used for regression.

| Term | Estimate | Std Error | t Ratio | Prob>\|t\| |
|---|---|---|---|---|
| Intercept | 11.5826367 | 0.44596741 | 25.97 | <.0001 |
| SCF | -0.5634655 | 0.39596685 | -1.42 | 0.1625 |
| Flt3L | -0.0394127 | 0.39596685 | -0.1 | 0.9212 |
| IL-3 | -0.0839281 | 0.39596685 | -0.21 | 0.8332 |
| IL-7 | 5.35939537 | 0.58504564 | 9.16 | <.0001 |
| TNFa | -0.4133108 | 0.39596685 | -1.04 | 0.3028 |
| CXCL12 | 0.10208908 | 0.39596685 | 0.26 | 0.7979 |
| Flt3L*IL-3 | -0.1256408 | 0.4600719 | -0.27 | 0.7862 |
| SCF*IL-7 | 1.20911486 | 0.4600719 | 2.63 | 0.0121 |
| TNFa*CXCL12 | -0.2820543 | 0.4600719 | -0.61 | 0.5433 |
| IL-7*IL-7 | -0.1251572 | 0.33575505 | -0.37 | 0.7113 |
| Block[1] | 3.0954772 | 0.49020658 | 6.31 | <.0001 |
| Block[2] | 0.6788045 | 0.4964669 | 1.37 | 0.1792 |
| IL-7*IL-7*IL-7 | -0.7262834 | 0.1964268 | -3.7 | 0.0007 |

**Supplemental Figure 26 (Related to Figure 4).** Linear regression model terms for CD4ISP cells from day 21-28. Provided as raw values at the default precision in JMP 14 software that was used for regression.

| Term | Estimate | Std Error | t Ratio | Prob>\|t\| |
|---|---|---|---|---|
| Intercept | 103.164804 | 2.38017518 | 43.34 | <.0001 |
| SCF | 15.857939 | 1.56543736 | 10.13 | <.0001 |
| IL-7 | 55.5635627 | 2.31295199 | 24.02 | <.0001 |
| TNFa | -3.6147244 | 1.56543736 | -2.31 | 0.0261 |
| CXCL12 | 0.71757033 | 1.56543736 | 0.46 | 0.6491 |
| SCF*IL-7 | 14.8698393 | 1.81887385 | 8.18 | <.0001 |
| TNFa*CXCL12 | -3.485291 | 1.81887385 | -1.92 | 0.0623 |
| SCF*SCF | -3.1972287 | 1.33007366 | -2.4 | 0.0208 |
| IL-7*IL-7 | -8.1344651 | 1.33007366 | -6.12 | <.0001 |
| TNFa*TNFa | 1.03273376 | 1.33007366 | 0.78 | 0.4419 |
| Block[1] | 9.88119172 | 1.93802454 | 5.1 | <.0001 |
| Block[2] | -0.4409151 | 1.96491797 | -0.22 | 0.8236 |
| IL-7*IL-7*IL-7 | -8.2384403 | 0.77656464 | -10.61 | <.0001 |

**Supplemental Figure 27 (Related to Figure 4).** Linear regression model terms for early DP cells from day 21-28. Provided as raw values at the default precision in JMP 14 software that was used for regression.

| Term | Estimate | Std Error | t Ratio | Prob>\|t\| |
|---|---|---|---|---|
| Intercept | 86.6349143 | 2.0179206 | 42.93 | <.0001 |
| SCF | 19.1067611 | 1.32718311 | 14.4 | <.0001 |
| IL-7 | 35.4637233 | 1.96092856 | 18.09 | <.0001 |
| TNFa | 1.84469728 | 1.32718311 | 1.39 | 0.172 |
| CXCL12 | 0.21150468 | 1.32718311 | 0.16 | 0.8742 |
| SCF*IL-7 | 12.3528251 | 1.54204743 | 8.01 | <.0001 |
| TNFa*CXCL12 | -2.3933864 | 1.54204743 | -1.55 | 0.1283 |
| SCF*SCF | -1.3848479 | 1.12764097 | -1.23 | 0.2264 |
| IL-7*IL-7 | -7.1153746 | 1.12764097 | -6.31 | <.0001 |
| TNFa*TNFa | 2.8558109 | 1.12764097 | 2.53 | 0.0152 |
| Block[1] | 10.9429091 | 1.64306379 | 6.66 | <.0001 |
| Block[2] | -0.8823612 | 1.66586413 | -0.53 | 0.5992 |
| IL-7*IL-7*IL-7 | -5.6418604 | 0.65837414 | -8.57 | <.0001 |

**Supplemental Figure 28 (Related to Figure 4).** Linear regression model terms for late DP cells from day 21-28. Provided as raw values at the default precision in JMP 14 software that was used for regression.

| Term | Estimate | Std Error | t Ratio | Prob>\|t\| |
|---|---|---|---|---|
| Intercept | 43.1526317 | 1.06544374 | 40.5 | <.0001 |
| SCF | 6.50420244 | 0.70074063 | 9.28 | <.0001 |
| IL-7 | 16.6457211 | 1.03535246 | 16.08 | <.0001 |
| TNFa | -0.9553494 | 0.70074063 | -1.36 | 0.1802 |
| CXCL12 | 0.19378692 | 0.70074063 | 0.28 | 0.7835 |
| SCF*IL-7 | 4.74860956 | 0.81418703 | 5.83 | <.0001 |
| TNFa*CXCL12 | -2.4891032 | 0.81418703 | -3.06 | 0.0039 |
| SCF*SCF | -1.2644009 | 0.59538419 | -2.12 | 0.0398 |
| IL-7*IL-7 | -3.3712565 | 0.59538419 | -5.66 | <.0001 |
| TNFa*TNFa | 1.81999508 | 0.59538419 | 3.06 | 0.0039 |
| Block[1] | 5.18271266 | 0.86752275 | 5.97 | <.0001 |
| Block[2] | -0.7477069 | 0.87956112 | -0.85 | 0.4002 |
| IL-7*IL-7*IL-7 | -2.6706032 | 0.34761556 | -7.68 | <.0001 |

**Supplemental Figure 29 (Related to Figure 4).** Linear regression model terms for CD8SP cells from day 21-28. Provided as raw values at the default precision in JMP 14 software that was used for regression.

| Term | Estimate | Std Error | t Ratio | Prob>\|t\| |
|---|---|---|---|---|
| Intercept | 32.2603708 | 0.7316126 | 44.09 | <.0001 |
| SCF | -0.2143526 | 0.48118042 | -0.45 | 0.6583 |
| IL-7 | 15.6124064 | 0.71094969 | 21.96 | <.0001 |
| TNFa | -1.1503109 | 0.48118042 | -2.39 | 0.0215 |
| CXCL12 | -0.0120103 | 0.48118042 | -0.02 | 0.9802 |
| SCF*IL-7 | -0.6638775 | 0.55908113 | -1.19 | 0.2419 |
| TNFa*CXCL12 | -0.8405346 | 0.55908113 | -1.5 | 0.1404 |
| SCF*SCF | -1.0381491 | 0.40883488 | -2.54 | 0.015 |
| IL-7*IL-7 | -2.4429343 | 0.40883488 | -5.98 | <.0001 |
| TNFa*TNFa | 0.68465812 | 0.40883488 | 1.67 | 0.1016 |
| Block[1] | 2.72608001 | 0.59570538 | 4.58 | <.0001 |
| Block[2] | -0.395117 | 0.60397182 | -0.65 | 0.5166 |
| IL-7*IL-7*IL-7 | -2.2515367 | 0.23869859 | -9.43 | <.0001 |

**Supplemental Figure 30 (Related to Figure 4).** Linear regression model terms for CD4ISP cells from day 28-35. Provided as raw values at the default precision in JMP 14 software that was used for regression.

| Term | Estimate | Std Error | t Ratio | Prob>\|t\| |
|---|---|---|---|---|
| Intercept | 53.5754183 | 0.75178221 | 71.26 | <.0001 |
| SCF | 9.32179642 | 0.73054966 | 12.76 | <.0001 |
| IL-7 | 4.50674811 | 0.73054966 | 6.17 | <.0001 |
| TNFa | -4.8876511 | 0.49444594 | -9.89 | <.0001 |
| SCF*IL-7 | 1.81056326 | 0.57449427 | 3.15 | 0.003 |
| IL-7*TNFa | -0.4952053 | 0.57449427 | -0.86 | 0.3937 |
| SCF*SCF | -1.3044505 | 0.42010593 | -3.11 | 0.0034 |
| IL-7*IL-7 | -2.846959 | 0.42010593 | -6.78 | <.0001 |
| TNFa*TNFa | -0.4528421 | 0.42010593 | -1.08 | 0.2874 |
| Block[1] | 1.32662297 | 0.61212821 | 2.17 | 0.0361 |
| Block[2] | 0.07712607 | 0.62062254 | 0.12 | 0.9017 |
| SCF*SCF*SCF | -0.8449201 | 0.24527921 | -3.44 | 0.0013 |
| IL-7*IL-7*IL-7 | 0.23670069 | 0.24527921 | 0.97 | 0.3402 |

**Supplemental Figure 31 (Related to Figure 4).** Linear regression model terms for early DP cells from day 28-35. Provided as raw values at the default precision in JMP 14 software that was used for regression.

| Term | Estimate | Std Error | t Ratio | Prob>\|t\| |
|---|---|---|---|---|
| Intercept | 71.5992903 | 0.99390129 | 72.04 | <.0001 |
| SCF | 15.5347015 | 0.96583058 | 16.08 | <.0001 |
| IL-7 | 3.66666305 | 0.96583058 | 3.8 | 0.0005 |
| TNFa | -7.6838777 | 0.65368727 | -11.75 | <.0001 |
| SCF*IL-7 | 1.65981404 | 0.75951598 | 2.19 | 0.0346 |
| IL-7*TNFa | -1.112947 | 0.75951598 | -1.47 | 0.1505 |
| SCF*SCF | -1.4944678 | 0.55540531 | -2.69 | 0.0103 |
| IL-7*IL-7 | -3.0759031 | 0.55540531 | -5.54 | <.0001 |
| TNFa*TNFa | -0.6824142 | 0.55540531 | -1.23 | 0.2262 |
| Block[1] | 2.23170456 | 0.80927031 | 2.76 | 0.0087 |
| Block[2] | 0.21447548 | 0.82050033 | 0.26 | 0.7951 |
| SCF*SCF*SCF | -1.7914292 | 0.32427386 | -5.52 | <.0001 |
| IL-7*IL-7*IL-7 | 0.79880273 | 0.32427386 | 2.46 | 0.0181 |

**Supplemental Figure 32 (Related to Figure 4).** Linear regression model terms for late DP cells from day 28-35. Provided as raw values at the default precision in JMP 14 software that was used for regression.

| Term | Estimate | Std Error | t Ratio | Prob>\|t\| |
|---|---|---|---|---|
| Intercept | 52.3310823 | 0.83554856 | 62.63 | <.0001 |
| SCF | 12.1459767 | 0.8119502 | 14.96 | <.0001 |
| IL-7 | 1.34598688 | 0.8119502 | 1.66 | 0.105 |
| TNFa | -4.615058 | 0.54953894 | -8.4 | <.0001 |
| SCF*IL-7 | 0.96339826 | 0.63850655 | 1.51 | 0.139 |
| IL-7*TNFa | -1.7068442 | 0.63850655 | -2.67 | 0.0107 |
| SCF*SCF | -0.9861832 | 0.46691569 | -2.11 | 0.0408 |
| IL-7*IL-7 | -1.6067212 | 0.46691569 | -3.44 | 0.0013 |
| TNFa*TNFa | -0.1688805 | 0.46691569 | -0.36 | 0.7194 |
| Block[1] | 2.82127714 | 0.6803338 | 4.15 | 0.0002 |
| Block[2] | 0.33549963 | 0.6897746 | 0.49 | 0.6293 |
| SCF*SCF*SCF | -1.1632524 | 0.27260912 | -4.27 | 0.0001 |
| IL-7*IL-7*IL-7 | 0.68930846 | 0.27260912 | 2.53 | 0.0154 |

**Supplemental Figure X (Related to Figure 4).** Linear regression model terms for CD8SP cells from day 28-35.

| Term | Estimate | Std Error | t Ratio | Prob>\|t\| |
|---|---|---|---|---|
| Intercept | 27.7659427 | 0.35325225 | 78.6 | <.0001 |
| SCF | 1.11467219 | 0.34327536 | 3.25 | 0.0023 |
| IL-7 | 1.14175554 | 0.34327536 | 3.33 | 0.0019 |
| TNFa | -2.3960274 | 0.23233343 | -10.31 | <.0001 |
| SCF*IL-7 | 0.04770922 | 0.26994705 | 0.18 | 0.8606 |
| IL-7*TNFa | -0.2525654 | 0.26994705 | -0.94 | 0.355 |
| SCF*SCF | -0.3045144 | 0.19740207 | -1.54 | 0.1306 |
| IL-7*IL-7 | -1.1775153 | 0.19740207 | -5.97 | <.0001 |
| TNFa*TNFa | -0.8254243 | 0.19740207 | -4.18 | 0.0001 |
| Block[1] | 0.52141125 | 0.28763073 | 1.81 | 0.0772 |
| Block[2] | -1.602835 | 0.2916221 | -5.5 | <.0001 |
| SCF*SCF*SCF | 0.18998604 | 0.11525337 | 1.65 | 0.1069 |
| IL-7*IL-7*IL-7 | 0.44170581 | 0.11525337 | 3.83 | 0.0004 |

**Supplemental Figure 33 (Related to Figure 4).** Linear regression model terms for CD4ISP cells from day 35-42. Provided as raw values at the default precision in JMP 14 software that was used for regression.

| Term | Estimate | Std Error | t Ratio | Prob>\|t\| |
|---|---|---|---|---|
| Intercept | 15.0784693 | 0.24322662 | 61.99 | <.0001 |
| SCF | 0.6406532 | 0.18301698 | 3.5 | 0.001 |
| IL-7 | 0.71577519 | 0.18301698 | 3.91 | 0.0003 |
| TNFa | -0.6432405 | 0.18301698 | -3.51 | 0.001 |
| IL-7*IL-7 | -0.3023616 | 0.15533652 | -1.95 | 0.0577 |
| TNFa*TNFa | -0.2681457 | 0.15533652 | -1.73 | 0.091 |
| Block[1] | 1.06565489 | 0.22657566 | 4.7 | <.0001 |
| Block[2] | 0.17154972 | 0.22958879 | 0.75 | 0.4587 |

**Supplemental Figure 34 (Related to Figure 4).** Linear regression model terms for early DP cells from day 35-42. Provided as raw values at the default precision in JMP 14 software that was used for regression.

| Term | Estimate | Std Error | t Ratio | Prob>\|t\| |
|---|---|---|---|---|
| Intercept | 35.9457562 | 0.67086437 | 53.58 | <.0001 |
| SCF | 6.26474675 | 0.50479497 | 12.41 | <.0001 |
| IL-7 | 2.92834805 | 0.50479497 | 5.8 | <.0001 |
| TNFa | -4.2569464 | 0.50479497 | -8.43 | <.0001 |
| IL-7*IL-7 | -1.4741273 | 0.42844711 | -3.44 | 0.0012 |
| TNFa*TNFa | -1.2646811 | 0.42844711 | -2.95 | 0.005 |
| Block[1] | 3.26100196 | 0.62493794 | 5.22 | <.0001 |
| Block[2] | 0.3823033 | 0.63324869 | 0.6 | 0.549 |

**Supplemental Figure 35 (Related to Figure 4).** Linear regression model terms for late DP cells from day 35-42. Provided as raw values at the default precision in JMP 14 software that was used for regression.

| Term | Estimate | Std Error | t Ratio | Prob>\|t\| |
|---|---|---|---|---|
| Intercept | 47.8080595 | 1.09075514 | 43.83 | <.0001 |
| SCF | 11.8192798 | 0.82074371 | 14.4 | <.0001 |
| IL-7 | 3.42460859 | 0.82074371 | 4.17 | 0.0001 |
| TNFa | -4.3786688 | 0.82074371 | -5.34 | <.0001 |
| IL-7*IL-7 | -1.7459706 | 0.69661008 | -2.51 | 0.0158 |
| TNFa*TNFa | -1.3026167 | 0.69661008 | -1.87 | 0.0679 |
| Block[1] | 6.59695715 | 1.01608358 | 6.49 | <.0001 |
| Block[2] | 1.85039497 | 1.02959599 | 1.8 | 0.0789 |

**Supplemental Figure 36 (Related to Figure 4).** Linear regression model terms for late DP cells from day 35-42. Provided as raw values at the default precision in JMP 14 software that was used for regression.

| Term | Estimate | Std Error | t Ratio | Prob>\|t\| |
|---|---|---|---|---|
| Intercept | 32.2984565 | 0.39517481 | 81.73 | <.0001 |
| SCF | 3.30658065 | 0.2973511 | 11.12 | <.0001 |
| IL-7 | 1.85584257 | 0.2973511 | 6.24 | <.0001 |
| TNFa | -2.9674504 | 0.2973511 | -9.98 | <.0001 |
| IL-7*IL-7 | -0.8202399 | 0.25237814 | -3.25 | 0.0022 |
| TNFa*TNFa | -0.4319561 | 0.25237814 | -1.71 | 0.0937 |
| Block[1] | 2.38903337 | 0.3681217 | 6.49 | <.0001 |
| Block[2] | 0.346024 | 0.37301717 | 0.93 | 0.3584 |

**Supplemental Figure 37 (Related to Figure 4).** Three-stage optimum cytokine concentrations (ng/ml).

| | Interval (days) | SCF | Flt3L | IL-3 | IL-7 | TNFα | CXCL12 |
|---|---|---|---|---|---|---|---|
| **Stage I** | 0-7 | 23.9 | 8.7 | 5.3 | 10.0 | 4.9 | 9.7 |
| **Stage II** | 7-21 | 120.5 | 8.0 | 1.2 | 44.5 | 0.4 | 14.8 |
| **Stage III** | 21-42 | 77.1 | 9.8 | 1.0 | 33.5 | 0.1 | 15.7 |

**Supplemental Table 38 (Related to Figures 1-3).** HSPC and proT-cell antibodies used in this study.

| Antigen | Fluorochrome | Clone | Cat. Num. | Vendor |
|---|---|---|---|---|
| CD117 (c-Kit) | AlexaFluor-488 | 104D2 | 313233 | Biolegend |
| CD123 (IL-3R) | BV421 | 9F5 | 562517 | BD Biosciences |
| CD127 (IL-7R) | PE Cy7 | A019D5 | 351320 | Biolegend |
| CD14 | FITC | M5E2 | 555397 | BD Biosciences |
| CD33 | FITC | HIM3-4 | 303304 | Biolegend |
| CD34 | PE | 581 | 555822 | BD Biosciences |
| CD34 | APC | 581 | 555824 | BD Biosciences |
| CD38 | PE | HIT2 | 555460 | BD Biosciences |
| CD5 | PE Cy7 | UCHT2 | 25-0059-42 | eBiosciences |
| CD56 | BV605 | NCAM16.2 | 562780 | BD Biosciences |
| CD7 | APC | M-T701 | 561604 | BD Biosciences |

**Supplemental Table 39 (Related to Figure 4).** T-cell maturation antibodies used in this study.

| Antigen | Fluorochrome | Clone | Cat. Num. | Vendor |
|---|---|---|---|---|
| CD3 | APC Cy7 | SK7 | 557832 | BD Biosciences |
| CD4 | BB515 | SK3 | 566912 | BD Biosciences |
| CD5 | PE Cy7 | UCHT2 | 25-0059-42 | eBiosciences |
| CD7 | APC | M-T701 | 561604 | BD Biosciences |
| CD8α | BV605 | SK1 | 564116 | BD Biosciences |
| CD8β | PE Cy7 | SIDI8BEE | 25-5273-42 | eBiosciences |

**Supplemental Table 40 (Related to Figure 2).** Primer sequences used for qPCR.

| Gene | Forward | Reverse |
|---|---|---|
| *ACTB1* | CATGTACGTTGCTATCCAGGC | CTCCTTAATGTCACGCACGAT |
| *BCL11b* | TCCAGCTACATTTGCACAACA | GCTCCAGGTAGATGCGGAAG |
| *CEBPA* | GGAGCTGAGATCCCGACA | TTCTAAGGACAGGCGTGGAG |
| *DTX1* | ATCGGAGAAGGCTCTACAGG | CGTCTGGCCTCCTTTCTAACT |
| *E2a* | CCGACTCCTACAGTGGGCTA | CGCTGACGTGTTCTCCTCG |
| *GATA3* | GTTGGCCTAAGGTGGTTGTG | ACAGGCTGCAGGAATAGGGA |
| *HES1* | CCTGTCATCCCCGTCTACAC | CACATGGAGTCCGCCGTAA |
| *NOTCH1* | GAGGCGTGGCAGACTATGC | CTTGTACTCCGTCAGCGTGA |
| *SPI1* | TGCAATGTCAAGGGAGGGGG | AAACCCTTCCATTTTGCACGC |
| *TCF7* | TGCACATGCAGCTATACCCAG | TGGTGGATTCTTGGTGCTTTTC |
